# BAG3 coordinates astrocytic proteostasis of Alzheimer’s disease-linked proteins via proteasome, autophagy, and retromer complex interactions

**DOI:** 10.1101/2025.09.20.677505

**Authors:** Zachary M. Augur, Garrett M. Fogo, Courtney R. Benoit, Gizem Terzioglu, Zachary R. Murphy, Mason R. Arbery, Natacha Comandante-Lou, Duc M. Duong, Nicholas T. Seyfried, Philip L. De Jager, Tracy L. Young-Pearse

## Abstract

**Background:** Bcl-2-associated athanogene 3 (BAG3) is a mediator of chaperone assisted selective autophagy, and in the brain, most highly expressed in astrocytes. However, its role in astrocytes remains poorly defined. Given the genetic and pathological links of BAG3 to proteostasis and neurodegenerative diseases, we investigated how BAG3 contributes to astrocyte function and Alzheimer’s disease (AD).

**Methods:** SnRNA-seq of the human brain determined cell type expression of BAG3. CRISPR/Cas9 gene editing in human iPSCs, followed by tandem mass tag-mass spectrometry and RNA-sequencing was performed to assess proteomic and transcriptomic changes following BAG3 loss. Co-immunoprecipitation of BAG3 in human astrocytes defined the interactome, with top interactors being validated by western blot (WB), AlphaFold modeling, and proximity ligation assays. In astrocytes, autophagic flux, lysosomal phenotypes, proteasome activity, and endocytic uptake were measured in BAG3 KO and BAG3 WT. Finally, BAG3 expression was assessed in postmortem AD brain by WB and snRNA-seq, and its functional relevance to amyloid-β (Aβ) degradation was tested in co-cultures of BAG3 KO iAs with familial AD neurons.

**Results:** In human brain and iPSC models, BAG3 was most highly expressed in astrocytes. Further, BAG3 loss caused greater proteomic disruption in astrocytes than in neurons. In the absence of BAG3, astrocytes showed reduced autophagy, diminished lysosome abundance and activity, and decreased proteasome function. To uncover molecular binding partners of BAG3 that might influence these phenotypes, we performed co-immunoprecipitation, revealing interactions with HSPB8 and other heat shock proteins, proteasome regulators (PSMD5, PSMF1), and the retromer component, VPS35. Integration of BAG3 KO transcriptomic and proteomic datasets pinpointed AD-relevant proteins under post-translational control of BAG3, which included GFAP, BIN1, and HSPB8. HSPB8 levels were markedly reduced in BAG3-deficient astrocytes with overexpression partially rescuing its levels. Loss of astrocytic BAG3 impaired Aβ clearance in co-culture with APP/PSEN1 mutant neurons, directly linking BAG3 to a disease-relevant astrocyte function. Finally, analysis of postmortem brain tissue revealed BAG3 marks a stress-responsive astrocyte subtype in the brain of aged individuals with AD.

**Conclusions:** BAG3 binds to key regulators of autophagy, proteasome activity, and retromer function to coordinate astrocyte proteostasis, lysosomal function, and Aβ clearance. These findings position BAG3 as a potential therapeutic target and coordinator of glial protein quality control in neurodegeneration.

## INTRODUCTION

Mutations in Bcl-2-associated athanogene 3 (BAG3) are linked to a spectrum of human diseases which are known to affect the heart, skeletal muscle, and nervous system [7, 13, 41, 43, 44, 49]. In the heart, BAG3 coding mutations give rise to cardiomyopathies, while in the brain they contribute to neuropathies [1, 39, 43]. BAG3 has also emerged in genome-wide association studies (GWAS) as a Parkinson’s disease (PD)-associated gene and has been increasingly investigated in the context of Alzheimer’s disease (AD) and tau proteostasis [14, 16, 29, 32, 44]. Nearly all studies into BAG3 functionality in the brain have been focused on neuronal cell types, which demonstrated that BAG3 facilitates the clearance of tau and α-synuclein through its well-established role in chaperone-assisted selective autophagy (CASA) [14, 29, 44, 45]. However, single-nucleus RNA sequencing of human brain tissue reveals that BAG3 expression is highest in astrocytes [2, 15, 41, 47]. Consistent with this, our data demonstrate that BAG3 loss has a more pronounced effect on the proteomic landscape of astrocytes than on neurons, highlighting an underexplored, potentially critical role for BAG3 in astrocyte biology and neurodegenerative disease mechanisms.

BAG3 contains multiple domains that mediate protein-protein interactions, supporting a scaffolding or chaperone-like role in multicomponent protein complexes. In line with this, cell-type specific interactomes have been reported for BAG3, which include not only canonical heat shock and autophagy-related proteins, but also many regulators of vesicle trafficking, cytoskeletal components, and membrane proteins [28]. Importantly, the BAG3 interactome appears to be highly context-dependent. Minimal overlap is observed between the cell types that have been studied (cardiomyocytes, cancer cell lines, neurons, and HEK cells), beyond the expected CASA and heat shock response machinery [9, 17, 22, 28, 33, 53]. This raises the possibility that BAG3 may coordinate ubiquitous cellular responses but also unique functions within each cell type, a feature that is especially relevant in long-lived, post-mitotic cells. Given the high expression of BAG3 in astrocytes, defining its unique interactome in this context could uncover novel mechanisms.

Despite the limited research of BAG3 in astrocytes, the few studies that have investigated BAG3 in this cell type have shown intriguing results. Seidel, *et. al.* used immunohistochemistry of human brain to show that in regions of neurodegeneration, astrocytes exhibited elevated levels of HSPB8 and modest increases in BAG3 [37]. Another group investigated the role of Bag3 in mouse astrocytes and neurodegeneration following its upregulation in Bmal1-deficient astrocytes [41]. *Bmal1* knockdown in mouse primary astrocytes increased Bag3 expression and enhanced uptake of α-synuclein fibrils, an effect that was reduced when Bag3 was silenced [41]. Notably, astrocyte-specific Bag3 overexpression was sufficient to promote α-synuclein clearance in vitro and reduce pathology in vivo [41]. Another study using a mouse model of traumatic brain injury (TBI) found BAG3 protein is expressed at higher levels in GFAP+ astrocytes in both mouse and human post-mortem brain tissue from TBI, AD, and AD with TBI individuals [44], and overexpression of BAG3 enhanced autophagic functioning in HEK293T cells [44]. These studies support the potential relevance of BAG3 in the functioning of astrocytes in AD.

In this study, we aimed to elucidate the molecular mechanisms by which BAG3 contributes to protein quality control in astrocytes and explore its potential as a therapeutic target for neurodegenerative diseases. Through unbiased analyses of genetically engineered iPSC-derived astrocytes, we identified BAG3 binding partners and elucidated the functional importance of BAG3 in the autophagy-lysosome pathway and the ubiquitin proteasome system. In addition, we discovered a novel interaction between BAG3 and the retromer complex, a hub for protein sorting and trafficking. Finally, we characterized BAG3 across brain tissue from individuals with AD and showed an important subtype of astrocytes within the brain that express high levels of BAG3 and related heat shock proteins, which we hypothesize are induced by Aβ accumulation and contribute to its clearance. Using human cellular models, we show that BAG3 is upregulated in astrocytes in response to AD-relevant stimuli and that loss of BAG3 in astrocytes results in impaired Aβ clearance.

## METHODS

### Cell Culture

#### Induced pluripotent stem cell lines

Human iPSC work was performed following IRB review and approval through MGB/BWH IRB (2016P000867 and 2015P001676). iPSCs were generated from cryopreserved peripheral blood mononuclear cell (PBMC) samples from autopsied participants from the ROS and MAP cohorts. iPSCs were generated using Sendai reprogramming method [23]. iPSCs undergo a rigorous quality procedure that includes a sterility check, mycoplasma testing, karyotyping, and pluripotency assays performed by the New York Stem Cell Foundation (NYSCF) [23]. iPSCs were maintained using StemFlex Medium (Thermo Fisher Scientific). All cell lines were routinely tested for mycoplasma using a PCR kit (MP0035-1KT) and STR profiling to prevent potential contamination or alteration to the cell lines [23, 30].

#### CRISPR/Cas9 editing to generate BAG3 WT, HET, and KO iPSCs

The following lines were chosen for CRISPR/Cas9 mutagenesis of *BAG3*: BR33 and BR24, two iPSC lines derived from non-cognitively impaired individuals (male and female, respectively) from the Religious Orders Study and Rush Memory and Aging Projects (ROSMAP) cohorts [23]. gRNA ‘GCAGCGATTCCGAACTGAGG’ was used to target exon 2 within the BAG3 sequence, which is conserved within relevant BAG3 transcript isoforms. The Zhang Lab CRISPR Design website (crispr.mit.edu) was used to generate guide RNAs (gRNAs) with minimal off-target effects. gRNAs were cloned into the pXPR-003 vector. hiPSCs were electroporated with the sgRNA-encoding plasmid and Cas9, cells were isolated for monoclonal selection and Sanger sequenced to determine *BAG3* mutations. For each targeted iPSC line, one unedited/wildtype clone and two mutant clones (HET and KO) were chosen for downstream analyses. IDT CRISPR Cas9 guide RNA design checker was used to check for any potential off-target effects [30].

#### Differentiation of iPSCs to induced astrocytes (iAs)

iPSC-derived astrocytes (iAs) were differentiated following previously published papers with minor modifications [6, 23–25]. iPSCs were plated at 95k cells/cm^2^ on growth factor reduced Matrigel (Corning #354230) coated plates prior to virus transduction. Then, iPSCs were transduced with three lentiviruses – Tet-O-SOX9-puro (Addgene plasmid #117269), Tet-O-NFIB-hygro (Addgene plasmid #117271), and FUdeltaGW-rtTA (Addgene plasmid #19780). The cells were then replated at 200k cells/cm^2^ using StemFlex Medium (Thermo Fisher Scientific) and ROCK inhibitor (10 mM at D0). The media was changed daily with Expansion Media (EM) from days 1 to 3, and gradually switched from EM to FGF media from day 4 to 7. On day 8, cells were dissociated using Accutase, and plated at 84k cells/cm^2^ using FGF media. Doxycycline (2.5 μg/mL, Sigma) was added from day 1 to the end of the differentiation, puromycin (1.25 mg/mL, Gibco) was added on days 3 and 4 of the differentiation, and hygromycin (100mg/mL, InvivoGen) was added from days 4 to 6 of the differentiation. From day 8 to the end of differentiation day 21, cells were cultured with maturation media and fed every 2–3 days.

<u>Induced astrocyte protocol media</u>:

- Expansion Media: DMEM/F12 (Thermo Fisher Scientific), 10% FBS, 1% N2 Supplement (Stemcell Technologies), 1% GlutaMAX (Life Technologies)
- FGF Media: Neurobasal media, 2% B27, 1% NEAA, 1% GlutaMAX, 1% FBS, 8 ng/mL FGF, 5 ng/mL CNTF, 10 ng/mL BMP4
- Maturation Media: 1:1 DMEM/F12 and neurobasal media, 1% N2, 1% GlutaMAX, 1% Sodium Pyruvate, 5 µg/mL N-N-acetyl cysteine, 5 ng/mL heparin-binding EGF-like GF, 10 ng/mL CNTF, 10 ng/mL BMP4, 500 µg/mL dbcAMP

#### Differentiation of iPSCs to induced neurons (iNs)

iPSC-derived neurons (iNs) were differentiated following a previously published paper with minor modifications [23, 31, 52]. iPSCs were plated at a density of 95k cells/cm^2^ on plates coated with growth factor reduced Matrigel one day prior to virus transduction (Corning #354230). Then, iPSCs were transduced with two lentiviruses – pTet-O-NGN2-puro (Addgene plasmid #52047, a gift from Marius Wernig) and FUdeltaGW-rtTA (Addgene plasmid #19780, a gift from Konrad Hochedlinger). The cells were then replated at 200k cells/cm^2^ using StemFlex Medium (Thermo Fisher Scientific) and ROCK inhibitor (10 mM at day 0). The media was changed to KSR media (day 1), 1:1 of KSR and N2B media (day 2) and N2B media (day 3). On day 4, cells were dissociated using accutase and plated at 50k cells/cm^2^ using iN D4 media (NBM media + 1:50 B27 + BDNF, GDNF, CNTF (10 ng/mL, Peprotech). Doxycycline (2 μg/mL, Sigma) was added from day 1 to the end of the differentiation, and puromycin (5 mg/mL, Gibco) was added from day 2 to the end of the differentiation. On day 3, B27 supplement (1:100, Life Technologies) was added. From day 4 to the end of the differentiation, cells were cultured in iN day 4 media and fed every 2–3 days.

<u>Induced neuron protocol media</u>:

- KSR media: Knockout DMEM, 15% KOSR, 1x MEM-NEAA, 55 mM beta-mercaptoethanol, 1x GlutaMAX (Life Technologies).
- N2B media: DMEM/F12, 1x GlutaMAX (Life Technologies), 1x N2 supplement B (Stemcell Technologies), 0.3% dextrose (D-(+)-glucose, Sigma).
- NBM media: Neurobasal medium, 0.5x MEM-NEAA, 1x GlutaMAX (Life Technologies), 0.3% dextrose (D-(+)-glucose, Sigma).

#### iPSC Neuron-Astrocyte co-cultures

iNs and iAs were differentiated to day 15 on 6 well plates, as described above. On day 15, iAs were dissociated with accutase and replated on top of relevant iN cultures (contact) at 47k cells/cm^2^ [30].

<u>iPSC Neuron-Astrocyte co-culture media</u>

- Maturation Media (1:1 DMEM/F12 and neurobasal media, 1% N2, 1% GlutaMAX, 1% Sodium Pyruvate, 5 µg/mL N-N-acetyl cysteine, 5ng/mL heparin-binding EGF-like GF, 10 ng/mL CNTF, 10 ng/mL BMP4, 500 µg/mL dbcAMP) + 1:50 B27 (Life Technologies) + 10 ng/mL BDNF (Peprotech) + 10 ng/mL GDNF (Peprotech) + 2.5 μg/mL doxycycline (Sigma).

### Cell culture treatments

For cell cultures receiving pharmacological treatment, 24 hours prior to harvest, cells were fed with fresh iA media containing the following compounds where applicable:

*Bortezomib:* Day 20 iAs were treated with 5 nM bortezomib as previously described [19]. The vehicle for this inhibitor was DMSO.

*Chloroquine:* Day 20 iAs were treated with 80 µM chloroquine. The vehicle for this treatment was water.

*Heat Shock:* Day 21 iAs were placed in an incubator set to 42°C with 5% CO_2_ for 1 hour. Cells were harvested in RIPA immediately following heat shock exposure.

### BAG3 Overexpression

For BAG3 overexpression, 4 plasmids were generated with VectorBuilder and used for experimentation. A control plasmid with an ORF stuffer (pRP[Exp]-EGFP-CAG>ORF_Stuffer), a plasmid containing a FLAG-tagged human BAG3 sequence under the CAG promoter (pRP[Exp]-EGFP-CAG>hBAG3[NM_004281.4]*/FLAG), a plasmid containing a FLAG-tagged human P209L mutant BAG3 sequence under the CAG promoter (pRP[Exp]-EGFP-CAG>hBAG3[NM_004281.4]*(P209L) /FLAG), and a plasmid containing a FLAG-tagged human E455K mutant BAG3 sequence under the CAG promoter (pRP[Exp]-EGFP-CAG>{hBAG3(E455K)/[NM_004281.4]*} /FLAG).

#### iA BAG3 plasmid overexpression

On day 18 of iA differentiation, cells were dissociated in accutase (diluted 1:3 in PBS) and transfected with one of the four plasmids described above using the Amaxa P3 Primary Cell 4D-Nucleofector Kit (Lonza, #V4XP-3024) at a ratio of 5 µg of plasmid per 1 million cells.

Nucleofected cells were plated in 24 well plates at 100k cells/well. 72 hours following transfection, conditioned media was collected, and cells were harvested as described elsewhere for the given assay being performed.

#### HEK293T BAG3 plasmid overexpression

The day before plasmid transfection, cells were passaged to a 6 well plate with a density of 2 million cells per well. 24 hours after passaging, cells were transfected using the relevant DNA plasmids along with the Lipofectamine^TM^ 3000 Reagent (Invitrogen), per the user guide. Briefly, a Lipofectamine 3000 reagent mastermix in Opti-MEM media was formulated by combining 125 µl of Opti-MEM and 3.75 µl of Lipofectamine^TM^ 3000 reagent for each well of a 6 well plate. Separately, a DNA mastermix was created by combining 125 µl of Opti-MEM with 2.5 µg of DNA plasmid and 5 µl of P3000 reagent per well. Next, we combined equal amounts (125 µl per well of a 6 well plate) of the Lipofectamine^TM^ 3000 reagent master mix and the DNA mastermix. The combined mastermixes were incubated for 15 minutes at room temperature. Then, for each well of a 6 well plate, we spiked in 250 µl of the combined mastermix. Cells were harvested 48 hours after transfection as described elsewhere for the given assay being performed.

### Western blotting (WB)

Cells were lysed with RIPA lysis buffer (Thermo Fisher Scientific #89900) with the protease inhibitor (Complete TM mini protease inhibitor, Roche) and phosphatase inhibitor (phosphoSTOP, Roche) added freshly before lysis. Cells were lysed for 30 minutes on ice before transferring lysates to microcentrifuge tubes. Cell debris was pelleted by centrifugation (15,000 x g) for 15 minutes at 4°C. Supernatant (cell lysate) was collected and stored at –20°C until use. Protein concentration in cell lysate samples was determined with the Pierce BCA Protein Assay kit when applicable (ThermoFisher, #23225). Cell lysates were prepared with 4X LI-COR loading buffer (Fisher Scientific, #NC9779096) and 2.5% β-mercaptoethanol, centrifuged, and incubated at 95°C for 5 minutes. Samples were resolved using Novex NuPAGE^TM^ 4-12% Bis-Tris gels (ThermoFisher, #NP0336BOX) and NuPAGE^TM^ 1X MOPS-SDS or MES-SDS running buffer (ThermoFisher, #NP0001). Gel electrophoresis was run at 200V for 50 minutes. SeeBlue Plus2 (ThermoFisher, #LC5925) pre-stained protein standard was used for evaluation of molecular weight. The gel was extracted and transferred to a nitrocellulose membrane by incubation with 20% methanol tris-glycine transfer buffer at 400 mA for two hours. The transferred blot was blocked with Odyssey blocking buffer (LI-COR, #927-50100) for 1 hour at room temperature with agitation and incubated with primary antibody (diluted in blocking buffer) overnight at 4°C with agitation. Blots were incubated with LI-COR secondary antibody diluted 1:10,000 in TBST for 1 hour at room temperature with agitation. Blots were washed twice (10 minutes per wash) with TBST and stored in 1X TBS until imaging. Blots were imaged on a LI-COR Odyssey machine and quantified using ImageStudio software.

#### Autophagic Flux

On Day 20 of iA differentiation, astrocytes were fed with media containing 80 µM chloroquine for 24 hours along with vehicle control (water). After 24 hours, samples were harvested as described above. LC3B has two bands, LC3B-I (top band) and LC3B-II (bottom band).

Autophagic flux was calculated by subtracting the amount of LC3B-II in the vehicle treated controls from the quantified amount of LC3B-II in the chloroquine treated cells.

#### Revert™ 700 Total Protein Stain

When normalizing to total protein, WBs were stained with REVERT 700 total protein stain per manufacturer’s guidelines. Briefly, before blocking or antibody exposure, blots were covered in REVERT Staining Solution for 5 minutes, washed with Wash Solution two times, and imaged on the LI-COR Odyssey machine. Following imaging, blots were stripped with Destaining Solution and then proceeded to blocking. The proceeding protocol is then performed as described above.

| Antibody | Source | Identifier |
| --- | --- | --- |
| Rabbit anti-BAG3 | Proteintech | Cat# 10599-1-AP |
| Mouse anti-GAPDH | Proteintech | Cat# 60004-1-Ig |
| Rabbit anti-GFAP | Abcam | Cat# ab7260 |
| Mouse anti-VIM | Millipore | Cat# CBL202 |
| Chicken anti-TUJ1 | Novus | Cat# nb100-1612 |
| Rabbit anti-TAU | Cell Signaling Technologies | Cat# 46687S |
| Mouse anti-CD44 | Abcam | Cat# ab254530 |
| Rabbit anti-S100B | Abcam | Cat# ab52642 |
| Mouse anti-Ubiquitin | Millipore | Cat# Mab1510-I-100UG |
| Rabbit anti-LAMP1 | Cell Signaling | Cat# 9091S |
| Rabbit anti-RAB5 | Cell Signaling | Cat# 3547S |
| Rabbit anti-HSPB8 | Proteintech | Cat# 15287-1-AP |
| Rabbit anti-BIN1 | Abcam | Cat# Ab182562 |
| Goat anti-VPS35 | Abcam | Cat# AB10099 |
| Mouse anti-VPS26A | Atlas Antibodies | Cat# AMAb90967 |
| Goat anti-PSMF1 | Novus | Cat# NB100-1178 |
| Rabbit anti-PSMD5 | Proteintech | Cat# 11469-1-AP |
| Rabbit anti-PSMB5 | Bethyl | Cat# A303-847A-M |
| Rabbit anti-HSF1 | Cell Signaling | Cat# 4356S |
| Rabbit anti-LC3B | Novus | Cat# NB100-2220 |
| Rabbit anti-DYKDDDDK Tag | Cell Signaling | Cat# 2368S |
| Mouse anti-M2 FLAG Tag | Millipore | Cat# F1804 |

### Immunocytochemistry

Day 21 cells were washed with PBS and then fixed with 4% paraformaldehyde (PFA, Sigma) for 15 minutes at room temperature. Cells were blocked in 2% donkey serum (Jackson ImmunoResearch Laboratories) and 0.3% Triton-X-100 (Sigma) in PBS for 1 hour at room temperature with agitation. Primary antibodies were diluted in fresh donkey serum blocking buffer and cells were incubated with primary antibody solution overnight at 4°C. Then, cells were washed with PBS three times, incubated with secondary antibodies for 1 hour at room temperature with agitation, and then washed with PBS three times. Cells were treated with DAPI stain (1:1000 dilution) for 10 minutes at room temperature with agitation, followed by a final wash to prepare for imaging. Images were taken on a Zeiss LSM710 Confocal or Andor Dragonfly 600 Spinning Disk Confocal.

#### LAMP1 and RAB5 imaging and quantification

LAMP1 (1:200 dilution) and RAB5 (1:100 dilution) ICC was performed as described above. Cells were plated at 100k cells per well of a 24-well plate with glass coverslips and imaged using the Zeiss LSM710 Confocal microscope. The quantification pipeline was created using CellProfiler. First, primary objects were identified, including the DAPI nuclei, CD44 positive cell areas, and the LAMP1 puncta. Next, an ‘object mask’ was created to delineate CD44 positive areas per cell by relating the nuclei to the CD44 ICC stain. The number and intensity of LAMP1 positive puncta were then calculated for each CD44 positive cell/nuclei.

### Proteasome activity assay

Proteasome activity assays were performed as previously described [19]. Briefly, astrocytes or HEK293T cells were washed with ice-cold DPBS on Day 21 (for iAs), followed by homogenization in 25 mM Hepes-KOH (pH 7.5), 5 mM MgCl2, 10% glycerol, 0.1% NP-40, 1 mM dithiothreitol (DTT), 1 mM adenosine triphosphate (ATP), 0.1 mM phenylmethylsulfonyl fluoride (PMSF), 1 mM NaF, followed by sonication in a water bath sonicator (B2500A-MT, VWR) for 20 times for 10 seconds each, with 30 second pauses between pulses. Protein lysates were centrifuged at 10,000 × g for 10 minutes at 4°C. Samples normalized for protein concentration (5-10 μg total protein) were loaded into a black-walled 96-well plate. Proteasome activity was monitored using Infinite 200 PRO plate reader (Tecan) with the i-control microplate reader software (Tecan). The reaction was started by adding 20 μM Suc-LLVY-amc (for chymotrypsin-like activity) in 50 mM Tris, 40 mM KCl, 5 mM MgCl2, 0.05 mg/mL Bovine Serum Albumin (BSA), 1 mM ATP, 1 mM DTT, 1 mM NaF at the zero-time point. For background detection, 2 μM epoxomicin was added to the reactions with Suc-LLVY-amc. The activity was assayed by measuring the fluorescence intensity for 2 hours at 37°C in 1-min intervals (excitation 380 nm; emission 460 nm).

### Live Cell Imaging

#### DQBSA

DQ Red BSA reagents were purchased from Invitrogen. A 1 mg/mL stock solution of DQ-BSA was prepared in PBS. iAs were grown on glass 24 well plates at a density of 100k cells/well. iAs were differentiated normally and on day 21, cells were washed once with PBS and exposed to 1 μg/mL of DQ-BSA diluted from the stock solution. DQ-BSA was added for 1 hour at 37°C while protected from light. After 1 hour, DQ-BSA treated cells were washed two times with fresh PBS and fresh iA media was added. Immediately after fresh media was added, the astrocytes were monitored every 1 hour using an Incucyte S3 Live-Cell Analysis System. Images were filtered in a blinded manner for oversaturation of DQ-BSA signal using the Incucyte. The amount of DQ-BSA signal was normalized to cell area using the light microscope on the Incucyte. For all analyses, 2 hour post-chase values were used.

### Flow Cytometry

Astrocytes were plated at 2 million cells per 10 cm plate and on day 21 were prepared for flow cytometry. 1 hour before harvest, astrocytes used for a negative control were pre-treated with cytochalisin D (1 uM, Sigma Aldrich C8273). Following pretreatment, cells were treated with either BSA-Alexa-488 for 1 hour (1 μg/mL in PBS, Invitrogen A13100) or pHrodo^TM^ Green Dextran, 10,000 MW for 20 minutes (50 μg/mL in DMSO, Invitrogen P35368). Following treatment, Zombie staining solution was prepared using PBS and 1:1000 Zombie violet dye (BioLegend, ref: 423113) and added to each plate. Plates were incubated at room temperature for 20 minutes in the dark. After this final incubation, cells were gently washed once with PBS and then were dissociated using accutase. Following dissociation, cells were moved to a conical tube and centrifuged at 500 g for 5 min. After this centrifugation step, the supernatant was discarded. 300 μl of FACS buffer was added, the cell pellet was resuspended, and the sample was transferred into flow cytometry tubes to be run on the BD LSRFortessa^TM^ Cell Analyzer.

### TMT proteomic and data analysis

#### Sample processing

iNs and iAs were plated in 6 well plates (1 million and 500k cells per well, respectively) and differentiated to day 21. Media was removed from the wells and cells were washed two times with ice-cold DPBS. Samples were processed as previously described [10, 24]. Cells were lysed in 250 μL of urea lysis buffer (8LM urea, 100LmM NaHPO4, pH 8.5) with the protease inhibitor (Complete TM mini protease inhibitor, Roche) and phosphatase inhibitor (phosphoSTOP, Roche) added freshly before the lysis. All homogenization was performed using a Bullet Blender (Next Advance) according to manufacturer protocols. Briefly, each tissue piece was added to urea lysis buffer in a 1.5 mL Rino tube (Next Advance) harboring 750 mg stainless steel beads (0.9–2 mm in diameter) and blended twice for 5 min intervals in the cold room (4°C). Protein supernatants were transferred to 1.5 mL Eppendorf tubes and sonicated (Sonic Dismembrator, Fisher Scientific) 3 times for 5Lseconds with 15Lsecond intervals of rest at 30% amplitude to disrupt nucleic acids and subsequently vortexed. Protein concentration was determined with BCA assay, and samples were frozen in aliquots at −80°C. Protein homogenates (50 μg) treated with 1 mM dithiothreitol (DTT) at 25°C for 30Lminutes, followed by 5 mM iodoacetamide (IAA) at 25°C for 30Lminutes in the dark. Protein mixture was digested overnight with 1:100 (w/w) lysyl endopeptidase (Wako) at room temperature. The samples were then diluted with 50LmM NH4HCO3 to a final concentration of less than 2LM urea and then and further digested overnight with 1:50 (w/w) trypsin (Promega) at 25°C. Resulting peptides were desalted with a Sep-Pak C18 column (Waters) and dried under vacuum.

#### Tandem mass tag (TMT) labeling

Peptides were reconstituted in 100μl of 100mM triethyl ammonium bicarbonate (TEAB) and labeling performed using TMTPro isobaric tags (Thermofisher Scientific, A44520) as previously described [18, 35]. Briefly, the TMT labeling reagents were equilibrated to room temperature, and anhydrous ACN (200 mL) was added to each reagent channel. Each channel was gently vortexed for 5 minutes, and then 20 mL from each TMT channel was transferred to the peptide solutions and allowed to incubate for 1 hour at room temperature. The reaction was quenched with 5% (v/v) hydroxylamine (5mL) (Pierce). All 16 channels were then combined and dried by SpeedVac (LabConco) to approximately 100 mL and diluted with 1 mL of 0.1% (v/v) TFA, then acidified to a final concentration of 1% (v/v) FA and 0.1% (v/v) TFA. Peptides were desalted with a 60 mg HLB plate (Waters). The eluates were then dried to completeness. High pH fractionation was performed essentially as described with slight modification [35]. Dried samples were re-suspended in high pH loading buffer (0.07% v/v NH4OH, 0.045% v/v FA, 2% v/v ACN) and loaded onto a Water’s BEH (2.1mm 3 150mm with 1.7 mm beads). An Thermo Vanquish UPLC system was used to carry out the fractionation. Solvent A consisted of 0.0175% (v/v) NH_4_OH, 0.01125% (v/v) FA, and 2% (v/v) ACN; solvent B consisted of 0.0175% (v/v) NH_4_OH, 0.01125% (v/v) FA, and 90% (v/v) ACN. The sample elution was performed over a 25 minute gradient with a flow rate of 0.6 mL/min with a gradient from 0 to 50% B. A total of 96 individual equal volume fractions were collected across the gradient and dried to completeness using a vacuum centrifugation.

#### Data processing protocol

All raw files were searched using Thermo’s Proteome Discoverer suite (version 2.4.1.15) with Sequest HT. The spectra were searched against a human Uniprot database downloaded August 2020 (86,395 target sequences). Search parameters included 10ppm precursor mass window, 0.05LDa product mass window, dynamic modifications methionine (+15.995LDa), deamidated asparagine and glutamine (+0.984LDa), phosphorylated serine, threonine, and tyrosine (+79.966LDa), and static modifications for carbamidomethyl cysteines (+57.021LDa) and N-terminal and Lysine-tagged TMT (+304.207LDa). Percolator was used to filter PSMs to 0.1%. Peptides were grouped using strict parsimony and only razor and unique peptides were used for protein level quantitation. Reporter ions were quantified from MS2 scans using an integration tolerance of 20 ppm with the most confident centroid setting. Abundance data were log_2_ transformed and batch effects were regressed using ComBat [26]. Differential expression was calculated using the DEP package in R [51].

### Gene set enrichment analysis (GSEA) and Gene Concept Network (GCN)

Gene set enrichment analysis, queried against the Gene Ontology database, was performed using the R package fgsea with a rank file derived from the summary statistics of the differential protein expression (–log_10_(p-value) * sign of the log_2_(FC)). Results were plotted using the R package ggplot2 (gene ratio = the number of leading-edge genes for a given pathway divided by the total size of that pathway). Genes were selected for pathway enrichment analysis by subsetting genes across three datasets: (1) BAG3 KO vs WT iA proteomic dataset, (2) ROSMAP iA Proteomic and RNAseq integration plots showing correlations of all proteins and genes to BAG3, respectively, and (3) correlations of BAG3 protein to all other proteins across 971 ROSMAP brain proteomics [38]. The cutoffs for each dataset are as follows: (1) q-value < 0.05 and log_2_(FC) > 0.2 or log_2_(FC) < –0.2, (2) ROSMAP BAG3 iA correlation for both RNAseq and proteomics is Pearson r > 0.4 or < –0.4 [25], (3) ROSMAP brain correlation of BAG3 to all other proteins is q-value < 0.05 and Pearson r > 0.2 or < –0.2 [38]. Overrepresentation analysis of the resulting gene lists was performed, queried against the Gene Ontology pathway database, and results were plotted using the R package ggplot2.

### Proteostasis Consortium chi squared testing

Significantly upregulated and downregulated proteins identified in the BAG3 WT vs. KO iA proteomic dataset were used to establish a baseline ratio of differentially expressed proteins. This baseline was then applied to subsequent chi-square analyses testing for enrichment within specific subsets of genes. Protein lists corresponding to distinct components of the autophagy-lysosome pathway (ALP) were curated using annotations from the Proteostasis Consortium (https://www.proteostasisconsortium.org). Separate chi-square tests were performed for each curated gene set to assess whether any specific arm of the ALP was disproportionately represented or suppressed among BAG3-regulated proteins.

### RNAseq

RNA was harvested using the PureLink RNA Mini Kit (ThermoFisher, #12183018A). Purity and concentration of RNA extractions were assessed using a NanoDrop spectrophotometer. Samples were sent to Genewiz (Azenta) for library preparation and sequencing. Fastq files were assessed using fastqc. Reads were pseudo-aligned using Kallisto and imported into Sleuth for differential expression analysis [3]. Counts were normalized, filtered for low abundance transcripts, and batch effects were regressed using ComBat [26]. Principal component analysis (PCA) was performed on the batch-corrected, log-transformed expression matrix using the R function prcomp. Differential expression was assessed using the Sleuth package in R [34].

Heatmaps were produced using the R package pheatmap with color palettes from the R package Seurat [5]. Gene set enrichment analysis, queried against the Gene Ontology database, was performed using the R package fgsea with a rank file derived from the summary statistics of the differential gene expression (–log_10_(p-value) * sign of the log_2_(FC)). Results were plotted using the R package ggplot2 (gene ratio = the number of leading-edge genes for a given pathway divided by the total size of that pathway).

### Immunoprecipitation followed by mass spectrometry (IP-MS)

BAG3 KO and WT iA pairs from both BR33 and BR24 were lysed in 1% CHAPSO buffer and assessed for protein concentration using the Pierce BCA Protein Assay kit (ThermoFisher, #23225). Lysates were pre-cleared with Protein G Dynabeads (Invitrogen, #10003D). Primary antibody was cross-linked to Protein G Dynabeads with BS_3_ (ThermoFisher, #21580). 5 μg of BAG3 primary antibody was used for immunoprecipitation. Immunoprecipitation was performed by incubating pre-cleared lysates with the cross-linked antibody-Dynabead complex overnight at 4°C. For immunoprecipitation followed by western blot (to validate specificity of antibody), beads were washed 3 times with CHAPSO lysis buffer before eluting protein with 5% β-mercaptoethanol and loading sample onto a Novex NuPAGE 4-12% Bis-Tris gel (ThermoFisher, #NP0336BOX). For immunoprecipitation followed by mass spectrometry, beads were washed with CHAPSO lysis buffer and frozen at –80°C. Samples were sent to the Emory Integrated Proteomics Core (Emory University) for peptide sequencing and identification. Enrichment analysis was performed with Perseus according to guidance from the Emory Integrated Proteomics Core.

### FLAG-Tag BAG3 Immunoprecipitation

FLAG-tagged immunoprecipitation was performed on HEK293T cells transfected with FLAG-tagged BAG3, as described above. Anti-FLAG M2 Magnetic Beads (Millipore, M8823) were used to immunoprecipitate BAG3 per the user guide. Briefly, 40 μl of the provided bead suspension was used per immunoprecipitation reaction within the relevant HEK293T cell lysate that was diluted to 0.1% CHAPSO. For each immunoprecipitation reaction, 1 well of a 6 well plate was used, where approximately 2 million cells per well were plated. A total of 1 mL diluted lysate was added to the M2 magnetic beads for each reaction. Samples and controls were nutated overnight at 4°C. The next day, samples were eluted in 2x sample buffer diluted in 0.1% CHAPSO and boiled for 4 minutes. The magnetic beads were separated from the eluted samples in sample buffer and then run on a western blot, as described previously.

### Alpha Fold Modeling

The full-length, unique protein sequence for each protein was obtained from UniProt and submitted to DeepMind’s AlphaFold Server with Seed set to ‘Auto’ (https://alphafoldserver.com/). Upon completion of the predicted alignment and folding, the AlphaFold 3 files were uploaded to the AlphaFold 3 Analysis Tool between June 1 and August 1, 2025 (https://predictomes.org/tools/af3/). Predicted alignment error (PAE) plots were generated from this pipeline for BAG3 and the following proteins: PSMF1, PSMD5, PSMB5, HSPB8, VPS35, VPS26A/B, and VPS29 [36].

### Proximity Ligation Assay

On Day 21, iA cultures were fixed with 4% PFA on glass coverslips. Coverslips were then permeabilized with 0.3% Triton-X 100 (Sigma, T8787) in DPBS for 60 minutes at room temperature. Coverslips were incubated in Duolink Blocking Solution (Millipore Sigma, DUO82007) for 60 minutes at 37°C. Coverslips were then incubated with primary antibodies for 1 hour at room temperature (1:300 Rb anti-FLAG and 1:100 Ms anti-VPS26A in Duolink Antibody Diluent [Millipore Sigma, DUO82008]). Coverslips were washed twice (5 min each) with Duolink Wash Buffer A (Millipore Sigma, DUO82049). Coverslips were incubated with secondary antibodies for 1 hour at 37°C (1:7 Duolink Ms PLUS [Millipore Sigma, DUO92001] and 1:7 Duolink Rb MINUS [Millipore Sigma, DUO92005] in Duolink Antibody Diluent [Millipore Sigma, DUO82008]). Coverslips were then washed twice (5 min each) with Duolink Wash Buffer A (Millipore Sigma, DUO82049). PLA probes were then ligated with incubation for 30 minutes at 37°C with 1:80 Duolink Ligase (Millipore Sigma, DUO82027) diluted in 1X Duolink Ligation Buffer (Millipore Sigma, DUO82009). Coverslips were then washed twice (5 minutes each) with Duolink Wash Buffer A (Millipore Sigma, DUO82049). Rolling circle amplification was performed during 30 min incubation at 37°C with 1:80 DNA polymerase (Millipore Sigma, DUO82028) diluted in 1X Amplification Buffer Red (Millipore Sigma, DUO82011). Coverslips were then washed twice (10 minutes each) with Duolink Wash Buffer B (Millipore Sigma, DUO82049).

Samples were then rinsed with 1:100 Wash Buffer B in nuclease-free H2O for 1 min. Coverslips were incubated with fluorescent secondary antibody (1:2000 anti-Rb Cy5 [Jackson Immunoresearch, 711-175-152]) for 1 hour at room temperature. Coverslips were then washed twice (10 min each) with Duolink Wash Buffer B (Millipore Sigma, DUO82049). Samples were then rinsed with 1:100 Wash Buffer B in nuclease-free H2O for 1 minute. Coverslips were mounted on glass slides using Fluoroshield with DAPI (Millipore Sigma, F6057). Imaging was performed on the Andor Dragonfly 600 Spinning Disk Confocal.

### Brain Extract Treatment

Human brain tissue applied to 9 iA genetic backgrounds was obtained from Albany Medical Center. Tris Buffered Saline (TBS: 20 mM Tris-HCl, 150 mM NaCl, pHL7.4) brain extracts were prepared using a method previously described [20, 40]. Briefly, the tissue was dissected to isolate the gray matter, which was homogenized in ice-cold TBS at a ratio of 1:4 tissue weight to buffer volume within a Dounce homogenizer. This suspension was then subject to ultra-centrifugation at 1.75L×L105 x g in a TL100 centrifuge (Beckman Coulter) to pellet cellular debris. The supernatant was placed in a 2-kDa dialysis cassette (Slide-A-Lyzer, Thermo Fisher) and dialyzed 1:104 (volume:volume) in artificial cerebrospinal fluid (aCSF: 124 mM NaCl, 2.5LmM KCl, 2.0LmM MgSO4, 1.25LmM KH2PO4, 26LmM NaHCO3, 10 mM glucose, 4LmM sucrose, 2.5LmM CaCl2). Following dialysis, the TBS brain extracts were aliquoted and stored at −L80°C. To treat iAs, the TBS brain extracts were ‘buffer-exchanged’ into culture media by placing extract in a 3-kDa centrifugation filter (Amicon Ultra, EMD Millipore) and performing a 90-min centrifugation at 3000 x g in an RC-5C centrifuge (Sorvall, Thermo Fisher). Following centrifugation, the retentate was reconstituted to 1X volume in BrainPhys Media with SM-1 neuronal supplement (StemCell Technologies). The protein concentration within the reconstituted extracts was determined by performing a bicinchoninic acid (BCA) assay (Thermo Fisher) on an 8-step, 2-fold dilution series of reconstituted extract and media to determine the proportion of protein contributed by the brain extract. The reconstituted extract was then back diluted in BrainPhys media to a concentration of 1 mg/mL brain extract protein. Day 18 iAs were treated with this human AD brain extract (BE) by diluting the 1 mg/mL BE 1:8 in iA culture media. Treatments were carried out for 72 hours at 37°C with media changed and fresh 1:8 AD BE added every 24 hours. After the final treatment, cells were washed three times with ice-cold DPBS and harvested in RIPA for WB, as described.

### ELISAs

48 hour conditioned media was collected immediately prior to harvest. Extracellular proteins were measured following manufacturer instructions for the Aβ Peptide Panel 1 (6E10) Kit. Samples were normalized to total protein concentration as determined by the Pierce BCA Protein Assay kit or Tuj1 measured in the lysates using WB (ThermoFisher, #23225).

### SnRNAseq Data Preprocessing and Subtype Identification in ROSMAP Brains

Processing and separation of cellular subpopulations was performed previously [15]. Briefly, data were derived from the dorsolateral prefrontal cortex of 437 randomly selected individuals in the ROSMAP cohort of patients. Nucleus isolation and single-nucleus RNA library preparation were performed in pooled batches of samples with 5,000 nuclei per individual being include for each sample. For each of the pooled batches the following was performed: (1) library alignment and background noise removal; (2) demultiplexing; (3) application of a normalization and clustering pipeline; (4) classification of nuclei for cell types; (5) removal of low-quality nuclei using a cell-type-specific threshold; and (6) detection of doublets for removal. Nuclei were then partitioned into subsets of different cell classes, and subclustering analysis was performed separately per cell type. After cleaning the nuclei per cell type, subclustering analysis per cell type was performed over multiple resolutions (Seurat FindClusters, algorithm=4, method=igraph), and the resolution was determined by the (1) differential gene expression per cluster (Seurat, FindAllMarkers, test. use=negbinom); (2) functional annotations of the differential signatures; and (3) the proportions of clusters across individuals. Clusters that did not have differential genes or clusters specific to a single individual were united with the neighboring cluster with a similar RNA profile. From here, BAG3 across astrocyte subclusters as well as analysis of leading edge genes in Ast. 9 subcluster was selected for additional analysis. To compare the gene expression across astrocyte subclusters, astrocyte raw gene counts were normalized in Seurat using the NormalizeData function (normalization.method = “LogNormalize”, scale.factor = 10000) followed by data scaling (ScaleData).

### Data visualization

Schematics were generated using Biorender. Graphs and heat maps were generated using R Studio or GraphPad Prism 10.

### Quantification and Statistical Analysis

Information regarding statistical analyses can be found in the figure legends. All statistical tests were performed using GraphPad Prism 10 or R Studio. All data is shown as mean ± SEM. Generally, comparisons between two groups were performed with student’s t test, and comparisons between more than two groups were analyzed using paired one-way ANOVA followed by Tukey’s or Sidak’s post-hoc testing. Pearson correlation method (linear relationship) was used for correlation analyses.

## RESULTS

### Astrocytes are exceptionally vulnerable to reductions in BAG3

While BAG3 has been extensively studied in the context of neurons, cardiomyocytes, and skeletal muscle, its function in other cell types is less understood. We first examined cell-type specific expression of BAG3 across the human brain. Using a previously published single-nucleus RNA sequencing (snRNA-seq) dataset derived from the dorsolateral prefrontal cortex (DL-PFC) of 437 participants in the Religious Orders Study and Rush Memory and Aging Projects (ROSMAP), we found that BAG3 is most abundantly expressed in astrocytes, with lower expression in other cell types (**Fig. 1A**) [15]. Due to its known role in proteostasis, BAG3 is especially critical in postmitotic cells. Thus, lower expression may not translate to a reduced importance in neurons. Considering the cell-type specific expression of BAG3, as well as the existing literature showing the importance of BAG3 in neurons, we aimed to interrogate and compare the consequences of BAG3 knockdown in human astrocytes and neurons.

**Figure 1.**
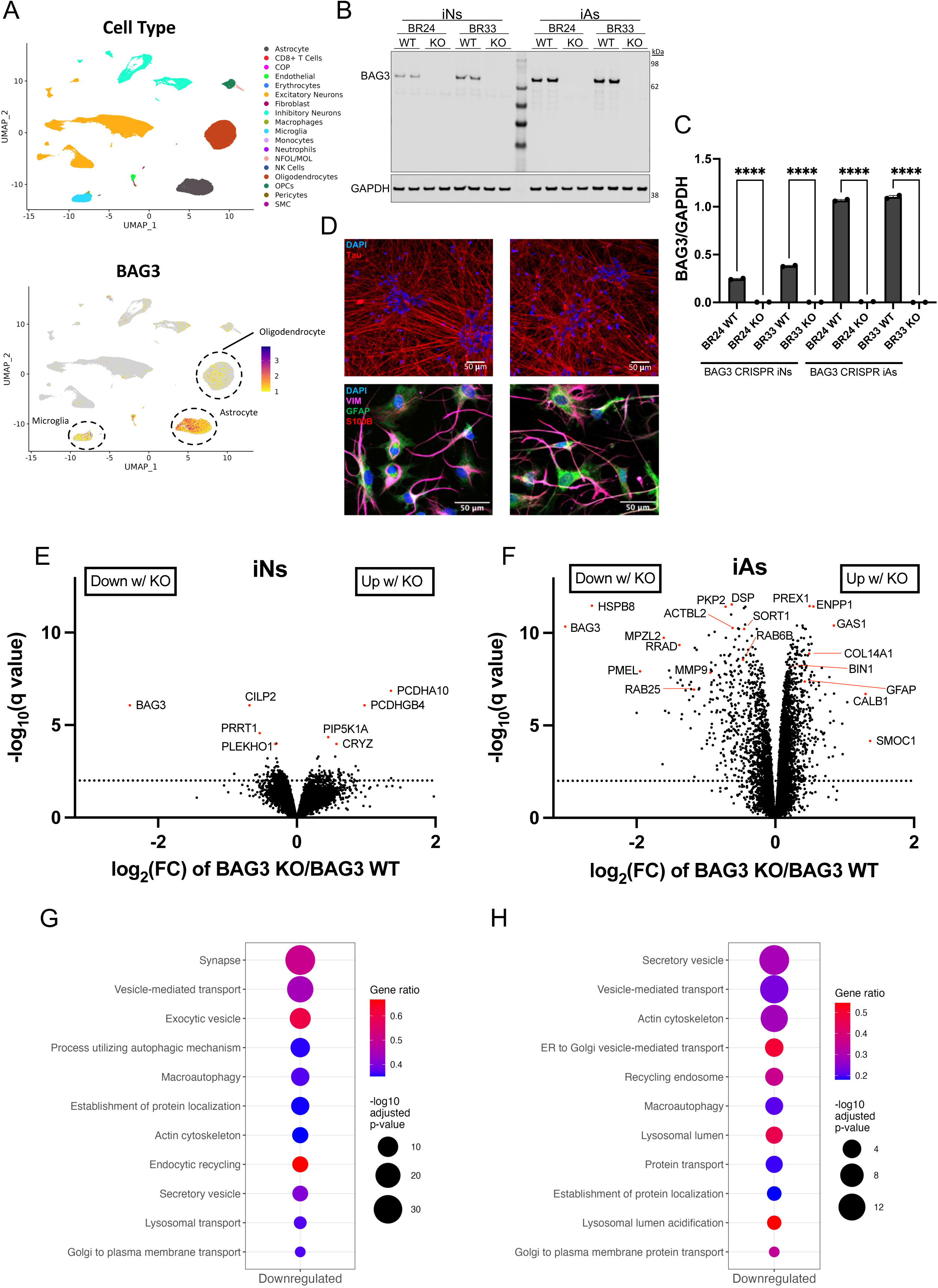
Proteome wide changes following BAG3 knockdown in astrocytes and neurons. (**A**) Single-nucleus RNA-sequencing (snRNAseq) of 437 ROSMAP dorsolateral prefrontal cortex samples showing the brain cell-types (top) [15]. BAG3 RNA expression across each cell-type is shown with astrocytes, oligodendrocytes, and microglia circled (bottom). (**B**) *BAG3* WT, HET, and KO iNs and iAs were produced using CRISPR/Cas9 technology targeting exon 2 of BAG3 in both BR24 and BR33 ROSMAP iPSCs. A representative western blot comparing iNs and iAs is shown. (**C**) The quantification of BAG3 knockdown in both iNs and iAs across both genetic backgrounds is shown (n = 2 independent wells, data shown as SEM, ANOVA with Tukey’s multiple comparison testing). (**D**) Immunocytochemistry for BAG3 WT and KO in both iNs and iAs are shown. iNs were stained with total tau antibody and DAPI. iAs were stained with DAPI, VIM, GFAP, and S100B antibodies. Scale bar = 50 µm (**E**) Volcano plot of BAG3 WT vs KO iNs showing differentially expressed proteins as measured by TMT-MS (dots above the dotted line; q < 0.01). Data shown includes BR24 and BR33 combined, with line ID regressed away (n = 3 wells per genetic background). (**F**) Volcano plot of BAG3 WT vs KO iAs showing differentially expressed proteins as measured by TMT-MS (dots above the dotted line; q < 0.01). Data shown includes BR24 and BR33 combined, with line ID regressed away (n = 4 wells per genetic background). Gene set enrichment analysis of the downregulated pathways with BAG3 KO using the Gene Ontology pathway database for (**G**) iNs and (**H**) iAs. For all comparisons: \**p* < 0.05, \*\**p* < 0.01, \*\*\**p* < 0.001, \*\*\*\**p* < 0.0001, ns = not significant.

To investigate the biological function of BAG3, we utilized CRISPR/Cas9 editing in two genetic backgrounds (one male, one female) to generate paired *BAG3* wild-type (WT) and knock-out (KO) induced pluripotent stem cell (iPSC) lines (**Supplementary Fig. S1**). These iPSCs were then differentiated into neurons (iNs) via NGN2 overexpression and into astrocytes (iAs) via SOX9 and NFIB overexpression, as previously described [6, 23, 25]. BAG3 KO was validated by western blot (WB) in both cell types (**Fig. 1B, Supplementary Fig S1C**). Consistent with expression in the brain, iAs expressed 4–5 times more BAG3 than iNs from the same genetic background, with efficient ablation of BAG3 protein observed in both cell types with KO (**Fig. 1C**). Differentiation efficiency was assessed by immunocytochemistry. iAs expressed GFAP, S100B, and VIM, and exhibited characteristic star-like morphology, while iNs expressed tau and showed characteristic neurite formation independent of BAG3 presence (**Fig. 1D**).

To compare proteome wide changes following BAG3 loss, we analyzed WT and KO iNs and iAs by TMT-MS. After filtering for protein detection, 10,330 proteins were quantified in iNs, with only 222 differentially expressed proteins (DEPs) consistently altered across both sets of lines (**Fig. 1E**). In contrast, 9,913 proteins were quantified in iAs, but 3,401 were consistently differentially expressed in response to BAG3 KO (**Fig. 1F**). Proteomic profiles confirmed efficient neuron and astrocyte differentiation across all lines (**Supplementary Table S1, S2**). Gene set enrichment analysis (GSEA) of DEPs against the Gene Ontology pathway database revealed downregulation of macroautophagy, transport pathways, actin cytoskeleton, and lysosomal pathways in both KO iNs (**Fig. 1G**) and iAs (**Fig. 1H**). To identify genes consistently associated with BAG3 expression, we calculated correlation coefficients between BAG3 and all genes measured at both RNA and protein levels across iAs from iPSCs from 43 ROSMAP participants using a previously published dataset (**Supplementary Fig. S2A-B**) [25]. This analysis revealed that in iAs, many BAG3-correlated genes are involved in protein transport and vesicular trafficking. A similar analysis was performed across 53 ROSMAP-derived iNs, which again identified several consistent genes and pathways (**Supplementary Fig. S2C-D**) [19]. To connect BAG3-dependent pathways in iAs to human brain tissue, we correlated BAG3 protein abundance with the abundance of all other proteins in 971 ROSMAP brain samples, and Gene Ontology (GO) pathway analysis was performed to identify coregulated pathways for BAG3 KO iAs, ROSMAP brain, and ROSMAP iAs (**Supplementary Fig. S2E**) [35]. Proteins downregulated in BAG3 KO iAs were used to construct a gene concept network that included proteins involved in actin filament binding, phospholipid binding, and peptidase regulator activity (**Supplementary Fig. S2F**). Strikingly, the recurrence of similar pathways and leading-edge genes across all datasets underscores the significance of these findings and the potential relevance of BAG3 to several biological functions in astrocytes including autophagy, protein/vesicular transport, and lysosomal function.

These unbiased analyses revealed a strong effect with loss of BAG3 and the findings encouraged further investigation into how BAG3 might be regulating some of these pathways and functions.

### The BAG3 interactome identifies autophagy, proteasome, and retromer component proteins as key binding partners in astrocytes

We next sought to understand the molecular mechanisms that may underlie the functional roles of BAG3 in astrocytes. To this end, we performed BAG3 immunoprecipitation in BAG3 WT and BAG3 KO iAs followed by data-independent acquisition mass spectrometry (DIA-MS) to define the human astrocyte BAG3 interactome. Differential protein abundance between WT and KO samples was used to identify high-confidence BAG3-binding partners (**Fig. 2A, Supplementary Table S3**). WB analysis of input and immunoprecipitated samples confirmed robust endogenous BAG3 pull-down in WT astrocytes and absence of BAG3 signal in KO controls (**Fig. 2B, Supplementary Fig. S3**). Known interacting partners of BAG3 such as HSPB8 and members of the HSPA family were identified, providing additional confidence for the identified novel interactors. In subsequent figures, we describe additional studies to validate these findings, with a focus on some of the strongest interactome hits: HSPB8, VPS35, and PSMD5/PSMF1.

**Figure 2.**
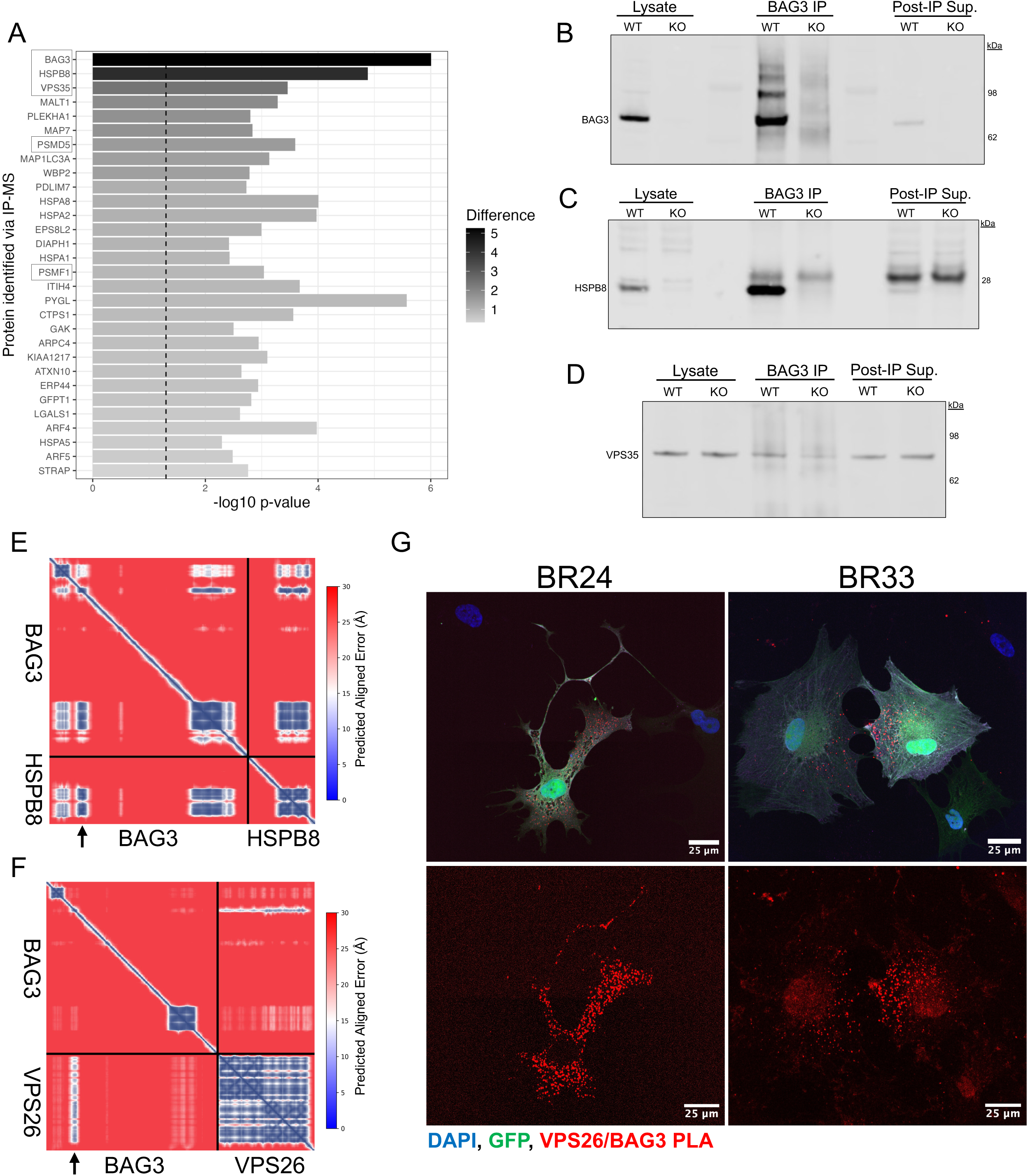
BAG3 astrocyte interactome implicates autophagy, ubiquitin proteasome system, and the retromer complex in BAG3 biological processes. (**A**) BR24 and BR33 WT and KO iAs were harvested in 1% CHAPSO lysis buffer and subjected to BAG3 antibody immunoprecipitation (IP) using Protein G Dynabeads. Samples were then sent for DIA-mass spectrometry. The top 30 most significantly enriched proteins in the BAG3 WT IP elution (p < 0.05) were graphed with proteins of interest boxed (n = 2 genetic backgrounds with 3 replicates each). (**B**) Representative western blot (WB) displaying the immunoprecipitation of BAG3 in WT and KO iAs using Protein G Dynabeads. (**C**) Representative WB displaying the co-immunoprecipitation of HSPB8 after immunoprecipitating for BAG3 in iAs. (**D**) Representative WB displaying the co-immunoprecipitation of VPS35 after immunoprecipitating for BAG3 in iAs. (**E-F**) Predicted aligned error (PAE) plot of BAG3 with (**E**) HSPB8 or (**F**) VPS26 using AlphaFold Multimer. Model 1 of 5 is shown with consistency observed across all 5 models. Arrows denote highest confidence predicted/known interacting sites between the two proteins. (**G**) Representative fluorescent images of a proximity ligation assay between FLAG-tagged BAG3 and VPS26. Red puncta represent positive signal or proximity between the two proteins. GFP positive cells represent positive BAG3 plasmid transduction. Representative images for both BR24 and BR33 are shown with only the red channel being shown separate from the whole image for ease of visualization.

As noted, HSPB8 has been shown to interact with BAG3 in other cell types and was completely ablated in our astrocyte TMT-MS dataset following loss of BAG3 [8]. Here, we validated this interaction in astrocytes through IP of BAG3 followed by WB of HSPB8 (**Fig. 2C, Supplementary Fig. S3**). Notably, VPS35, a core component of the retromer complex involved in endosomal sorting and trafficking, was the second strongest hit, and this was validated as a novel astrocyte BAG3 interactor by co-IP followed by WB in astrocytes (**Fig. 2D, Supplementary Fig. S3**). The predicted alignment error (PAE) plot derived from AlphaFold showed a robust predicted interaction between HSPB8 and BAG3, primarily around the isoleucine-proline-valine (IPV) domain of BAG3 (arrow) (**Fig. 2E**). To investigate the novel interaction of BAG3 and VPS35 further, we again performed AlphaFold-based protein–protein interaction modeling between BAG3 and each of the three core components of the retromer complex: VPS26, VPS29, and VPS35. The resulting PAE plots revealed the strongest and most confident interaction interface specifically between BAG3 and VPS26, with the highest-confidence contact centered around amino acid 208 of BAG3 (arrow), immediately adjacent to the site of the disease-associated P209L mutation (**Fig. 2F**). VPS35 and VPS29, the other two components of the retromer complex, also showed predicted interaction interfaces, but these had slightly lower confidence than VPS26 (**Supplementary Fig. S3A-B**). This finding aligns with our biochemical pulldown data and raises the possibility that BAG3 may physically associate with the entire retromer complex via this region of VPS26 or with other potential interacting sites in VPS29 or VPS35. To further validate this predicted interaction, we performed proximity ligation assays (PLA) in astrocytes overexpressing FLAG-tagged human BAG3 and probed for an interaction with endogenous retromer complex components. While available antibodies to BAG3 were specific via WB, these same antibodies when used for immunostaining showed robust signal in our BAG3 KO iAs. Thus, it was necessary to tag BAG3 for PLA experimentation. Given the PAE plot findings and the antibodies available for ICC, we selected VPS26 for PLA analysis. We observed robust PLA signal (red) in cells expressing FLAG-tagged BAG3 WT, whereas the empty vector control showed little to no signal (**Fig. 2G, Supplementary Fig. S3C**), supporting a direct or very close (within 40 nm) spatial interaction between BAG3 and the retromer complex in situ. While co-IP and PLA of two distinct retromer components support BAG3’s association with the full complex, we cannot fully exclude the possibility of an indirect interaction.

To extend these BAG3 interactome findings, we performed overexpression experiments in human embryonic kidney (HEK293FT) cells. We overexpressed FLAG-tagged human WT BAG3, two disease-associated BAG3 mutants (P209L and E455K), or an empty vector control. FLAG immunoprecipitation followed by WB for BAG3 confirmed robust and specific pulldown across all constructs, with the exception of P209L, along with no detectable BAG3 signal observed in the vector control (**Fig. 3A, Supplementary Fig. S4**). P209L mutants were used in this experimental setting, but due to protein instability and/or inefficient transduction at all DNA concentrations tested, results with this mutant are inconclusive. After validating this HEK293FT BAG3 OE system, we observed efficient co-immunoprecipitation of VPS35 with WT BAG3 as well as the mutant construct (**Fig. 3A, Supplementary Fig. S4**), supporting the specificity and reproducibility of this interaction. Next, we interrogated proteasome-related proteins as additional binding partners relevant in proteostasis. PSMD5 binding was validated in HEK cells overexpressing WT BAG3, but this interaction was ablated with E455K mutation (**Figure 3A, Supplementary Fig. S4**). PSMF1, another regulatory particle of the proteasome, was similarly validated in HEK cells but interestingly was not differentially immunoprecipitated with E455K mutation (**Fig. 3A, Supplementary Fig. S4**). To reinforce the specificity of these interactions to the regulatory particle of the proteasome, we also probed PSMB5, a component of the proteasome core particle, which showed no specific interaction with HEK cells overexpressing BAG3 (**Fig. 3A, Supplementary Fig. S4**). We then modeled the predicted BAG3 interaction with PSMF1, PSMD5, or PSMB5 using AlphaFold-based protein–protein interaction modeling which provided minimal insight into the binding dynamics with BAG3, emphasizing the importance of biochemical validation in conjunction with any AlphaFold based predictions (**Fig. 3 B-C, Supplementary Fig. S3D**).

**Figure 3.**
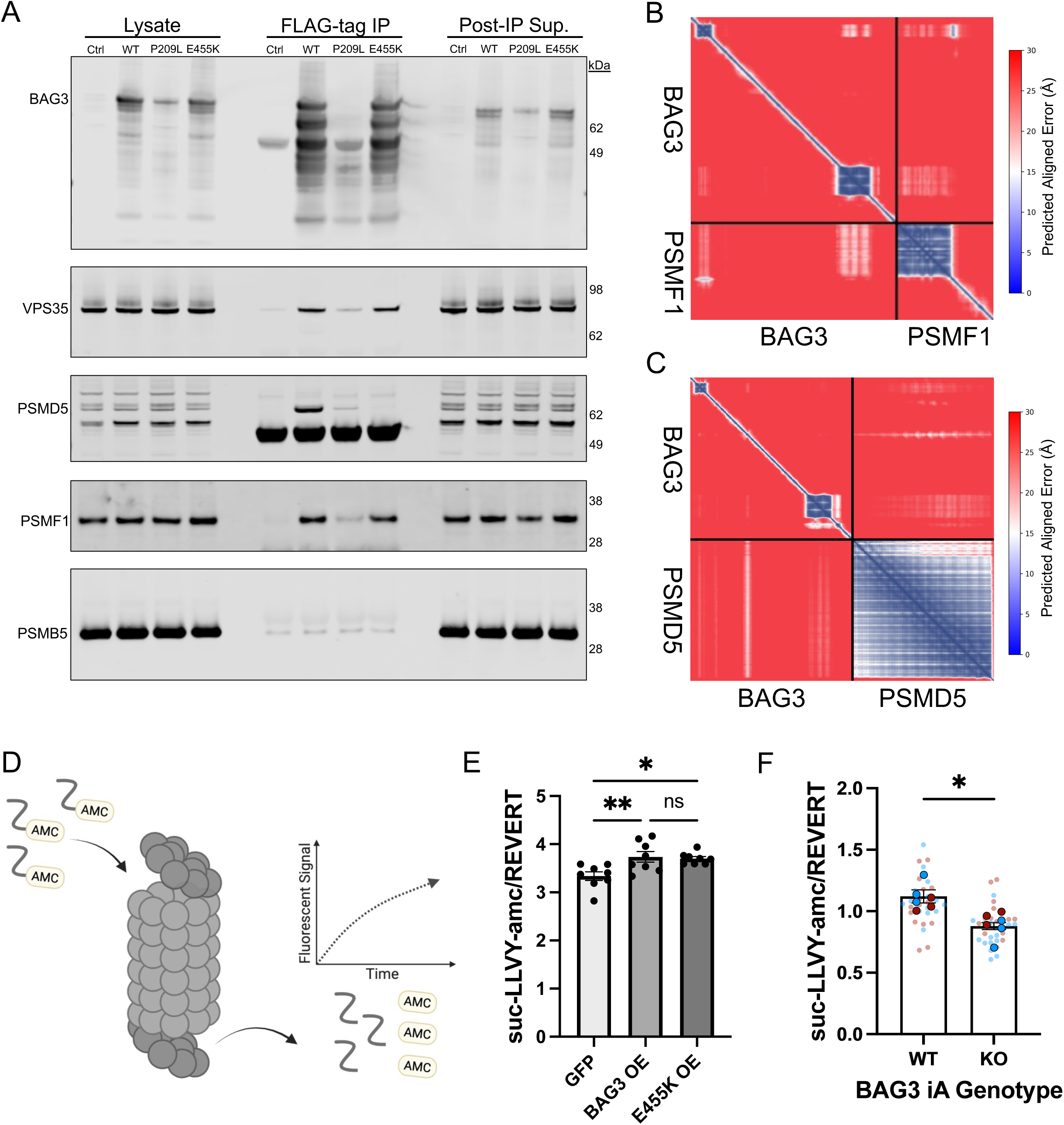
The BAG3 interactome validates in HEK293FT cells and its overexpression affects ubiquitin proteasome system (UPS) activity. (**A**) Representative WBs of FLAG-tagged BAG3 immunoprecipitation along with co-immunoprecipitation of VPS35, PSMD5, PSMF1, and PSMB5, in HEK293FT cells overexpressing either FLAG-tagged BAG3 WT, mutant P209L BAG3, mutant E455K BAG3, or empty vector control. (**B-C**) Predicted aligned error (PAE) plot of BAG3 with (**B**) PSMF1 or (**C**) PSMD5 using AlphaFold Multimer. Model 1 of 5 is shown with consistency observed across all 5 models. (**D**) Representative graphic showing the mechanism for the suc-LLVY-amc fluorogenic substrate, which measures chymotrypsin-like activity of the proteasome. (**E**) Quantification of the chymotrypsin-like activity of the proteasome in HEK293FT cells overexpressing either FLAG-tagged BAG3 WT, mutant E455K BAG3, or empty vector control. Data presented as slope of change from 20 minutes to 60 minutes following substrate addition (n = 8 replicates across 2 experiments; one-way ANOVA followed by Tukey’s multiple comparison testing). (**F**) Quantification of the chymotrypsin-like activity of the proteasome in BAG3 WT and KO iAs. Data presented as slope of change from 20 minutes to 60 minutes following substrate addition (n = 6 differentiations across 2 genetic backgrounds and 4 technical replicates per differentiation; paired t-test). Each solid dot corresponds to a single differentiation with technical replicates (transparent smaller dots) averaged. Blue = BR24, Red = BR33. Data presented with SEM. For all comparisons: \**p* < 0.05, \*\**p* < 0.01, \*\*\**p* < 0.001, \*\*\*\**p* < 0.0001, ns = not significant.

Next, taking advantage of our BAG3 OE model in HEK cells, we assessed proteasome activity to determine if BAG3-PSMD5/PSMF1 interactions were associated with functional consequences. Using the suc-LLVY-amc substrate to measure chymotrypsin-like activity in HEK cells overexpressing BAG3 WT, E455K mutant, or control, we saw an overall increase in proteasome activity with higher BAG3 (**Fig. 3D-E**). Despite loss of interaction with PSMD5 in the co-IP, no difference in elevation of proteosome activity was observed for the E455K mutant OE (**Fig. 3E**). Considering the observed increase in proteasome activity in HEK293FT cells overexpressing BAG3, we wanted to determine if loss of function of BAG3 would have the opposite effect in our BAG3 KO astrocyte model system. Here, we again measured proteasome activity using the suc-LLVY-amc substrate in our KO astrocytes and detected an overall reduction in proteasome activity with loss of BAG3 (**Fig. 3F**).

### BAG3 knockdown diminishes protein degradation and endo-lysosomal functioning

Given the strong interaction of BAG3 and HSPB8 in human astrocytes (along with many other autophagy-related proteins), the protein degradation changes in HEK cells overexpressing BAG3, and the established connection of BAG3 and the autophagy-lysosomal pathway (ALP) in other cell types, we returned to the proteomic profiling results of BAG3 KO iAs and investigated further the ALP-related protein changes observed in Fig. 1H. Using the Proteostasis Consortium’s Human Proteostasis Network ALP annotations, we categorized ALP components into sub-branches including lysosomal fusion, catabolism, and autophagy gene expression. In KO iAs, proteins relevant in lysosomal catabolism and fusion were decreased, while proteins used in autophagy substrate selection were elevated (**Fig. 4A**).

**Figure 4.**
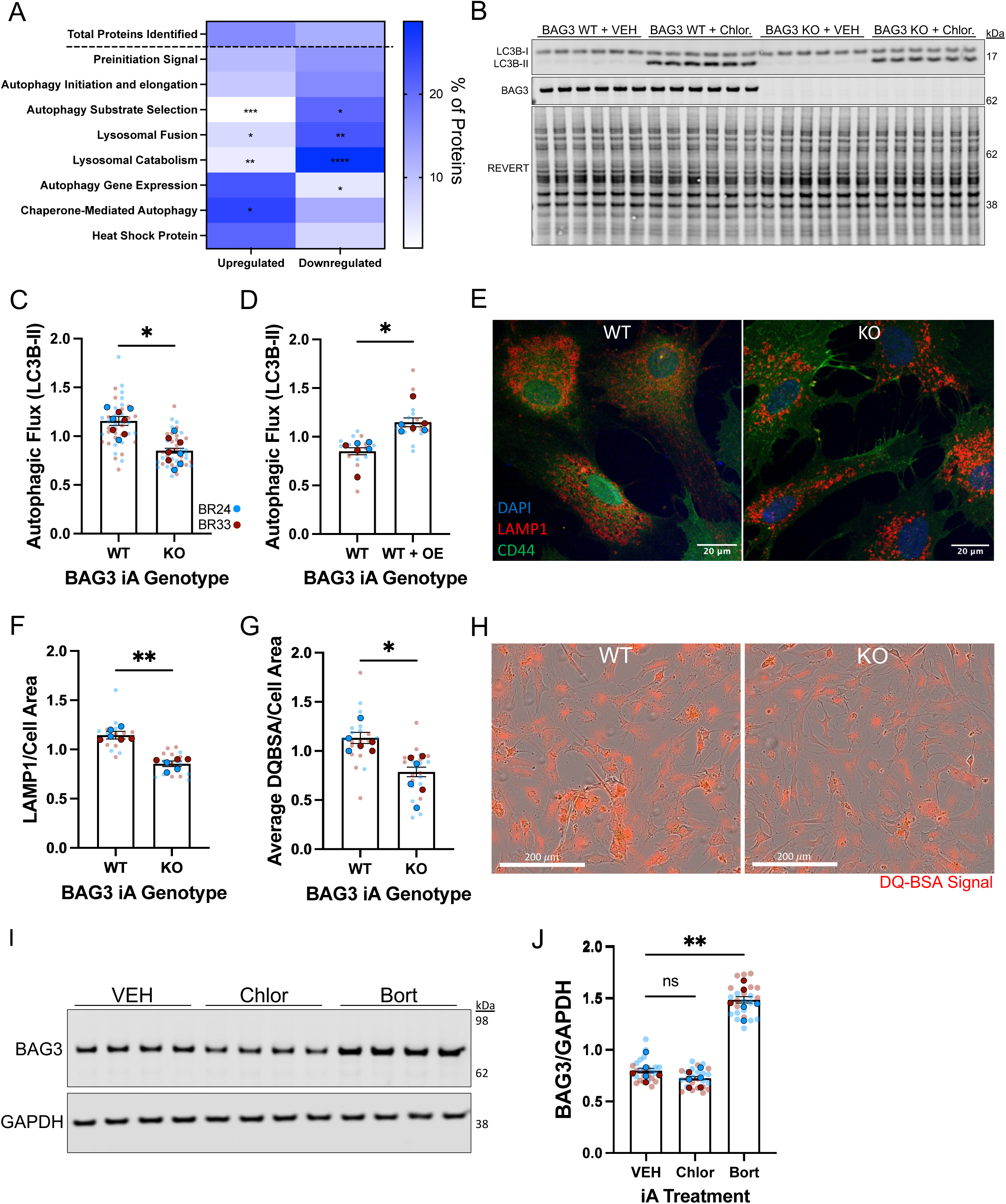
BAG3 knockdown in astrocytes diminishes protein degradation capacity and lysosomal functioning. (**A**) Autophagy lysosome pathway (ALP) proteins from the BAG3 WT vs KO iA TMT-MS (Figure 1F) were separated using the ALP terms derived from the Northwestern Proteostasis Consortium. Significance for each subset of genes was determined using chi squared testing against the total protein level change observed following BAG3 KO in iAs. The data is presented as the % of protein within that given ALP term. (**B**) *BAG3* WT and KO iA autophagic flux was measured after 24 hours of 80 µM chloroquine treatment, as assayed by LC3B-II accumulation in chloroquine treated samples compared to vehicle. Representative western blot (WB) is shown. (**C**) Quantification of autophagic flux following chloroquine treatment in BAG3 WT and KO iAs, normalized to REVERT (n = 8 differentiations across 2 genetic backgrounds and 3-6 technical replicates per differentiation). (**D**) Quantification of autophagic flux following 3 day BAG3 overexpression in WT iAs, normalized to GAPDH (n = 6 differentiations across 2 genetic backgrounds and 3 technical replicates per differentiation). (**E**) Representative LAMP1 staining with CD44 and DAPI in BAG3 WT and KO iAs used for quantification. Scale bar = 20 µm. (**F**) LAMP1 puncta per astrocyte cell area, as defined by CD44 staining, was quantified using cell-profiler (n = 6 differentiations across 2 genetic backgrounds and 3 technical replicates per differentiation, which includes 8 images per technical replicate). (**G**) Average DQ-BSA signal normalized to total cell area, measured 2 hours after DQ-BSA chase (n = 6 differentiations across 2 genetic backgrounds and 3 technical replicates per differentiation, which includes 9 images per technical replicate). (**H**) Representative phase contrast and florescent DQ-BSA Red signal captured at 2 hours post DQ-BSA chase using an Incucyte imaging system. Scale bar = 200 µm. (**I**) Representative WB showing BAG3 abundance following bortezomib or chloroquine treatment. (**J**) WB quantification of BAG3 normalized to GAPDH following chloroquine (80 µM for 24 hours) or bortezomib (5 nM for 24 hours) treatment (n = 6 differentiations across 2 genetic backgrounds, and 3-4 technical replicates per differentiation; paired one-way ANOVA with Tukey’s multiple comparison testing). Unless otherwise specified, statistics were performed using a paired t-test (2C-D, F-G) and all data is displayed using SEM. Each solid dot corresponds to a single differentiation with technical replicates (transparent smaller dots) averaged. Blue = BR24, Red = BR33. For all comparisons: \**p* < 0.05, \*\**p* < 0.01, \*\*\**p* < 0.001, \*\*\*\**p* < 0.0001, ns = not significant.

The pronounced dysregulation of ALP proteins in BAG3 KO astrocytes led us to measure autophagic flux to determine the functional impact of BAG3 loss in astrocytes (**Fig. 4B, Supplementary Fig. S5**). Consistent with proteomic profiling results, BAG3 KO iAs exhibited a significant reduction in autophagic flux, as measured by LC3B-II following chloroquine treatment (**Fig. 4C**). To further strengthen the relevance of BAG3 to astrocyte autophagy, we overexpressed BAG3 in the same genetic backgrounds and observed a corresponding increase in autophagic flux (**Fig. 4D**). Considering the significant reduction in autophagy following BAG3 ablation, we next investigated whether alterations in lysosomal function or abundance could underlie this phenotype. Quantification of lysosomal-associated membrane protein 1 (LAMP1), a key lysosomal glycoprotein, was performed using immunocytochemistry (ICC) in iAs. Total astrocyte area was marked with CD44 antibody, and the number of LAMP1-positive puncta were quantified per cell. This analysis revealed a reduction in total LAMP1-positive puncta in BAG3 KO iAs (**Fig. 4E-F**). To determine whether endosomal compartments also were affected, RAB5, an early endosome marker, was quantified, but showed no difference between WT and KO iAs (**Supplementary Fig. S5A-B**). Then, using a self-quenching DQ-BSA substrate, a significant reduction in lysosomal activity in BAG3 KO iAs was observed (**Fig. 4G-H**). To investigate whether this decrease was attributed to impaired endocytosis rather than lysosome dysfunction, we measured uptake of pHrodo Green Dextran and separately, constitutively fluorescent BSA-GFP, by flow cytometry. BAG3 KO iAs showed a modest but nonsignificant reduction in signal from both substrates (**Supplementary Fig. S5C-E**). These data suggest the reduction in DQ-BSA signal cannot be attributed solely to defects in endocytosis but is more likely due to a decrease in the total number of lysosomes or reduced enzymatic activity per lysosome following BAG3 KO.

Lastly, we investigated whether the reverse was true: could pharmacologic inhibition of autophagy or the proteasome alter BAG3 protein expression? iAs were treated with chloroquine (80 µM, 24 hours) to inhibit autophagy or bortezomib (5 nM, 24 hours) to inhibit the proteasome. WB analysis revealed BAG3 protein levels trended downward with autophagy inhibition and significantly increased following proteasome inhibition (**Figure 4I-J, Supplementary Fig. S5G**).

### Integration of transcriptomic and proteomic profiling data following BAG3 loss reveals post-transcriptional control of proteins relevant in protein degradation and neurodegeneration

We next aimed to analyze the effect of BAG3 disruption on the transcriptome of iAs using bulk RNA-sequencing to understand what protein level changes in BAG3 KOs are due to transcriptomic alterations rather than more direct defects in proteostasis. RNAseq was performed on WT and KO pairs of astrocytes in both genetic backgrounds (**Supplementary Table S4**). Gene-level differential expression of KO vs. WT iAs was calculated, and a volcano plot was used to visualize the results (**Fig. 5A, Supplementary Fig. S6A**). The top 25 upregulated (**Fig. 5B**) and downregulated (**Fig. 5C**) genes were visualized in a heat map showing significant concordance between genetic backgrounds. Pathway enrichment analysis was performed using the differentially expressed genes in BAG3 KO iAs queried against the Gene Ontology pathway database. This analysis showed downregulation of signaling receptor regulator activity, secretory vesicles, inflammatory response, proteolysis, and response to amyloid beta (**Fig. 5D**) with upregulation showing pathways involved in cholesterol biosynthesis (**Supplementary Fig. S6B**). Using the RNAseq and proteomic data sets, we created an integration plot of the DEGs/DEPs in BAG3 KO iAs (**Fig. 5E**). Genes in the top left (blue) quadrant, such as BIN1 and GFAP, were upregulated in BAG3 KO at the protein level compared to BAG3 WT, but unchanged or downregulated with RNA (**Fig. 5F**). Genes in the top right (green) quadrant were upregulated at both the protein and RNA level, such as FUS (**Supplementary Fig. S6C**). Genes in the bottom left (gray) quadrant were downregulated at both the protein and RNA level, such as ACTBL2 (**Supplementary Fig. S6C**). Finally, genes in the bottom right (red) quadrant were downregulated at the protein level, but not downregulated at the RNA level, suggesting some of these proteins could be stabilized by BAG3, such as HSPB8 (**Fig. 5F**). Of note, BAG3 knockdown does not consistently alter HSF1 nuclear localization or overall abundance following proteotoxic stress induced by heat shock or bortezomib exposure. As HSF1 is a master transcriptional regulator of the heat shock response, this suggests that the effect of BAG3 on downstream protein stability, such as HSPB8 loss, are unlikely to be mediated through impaired HSF1 activation (**Supplementary Fig. S6D-F**).

**Figure 5.**
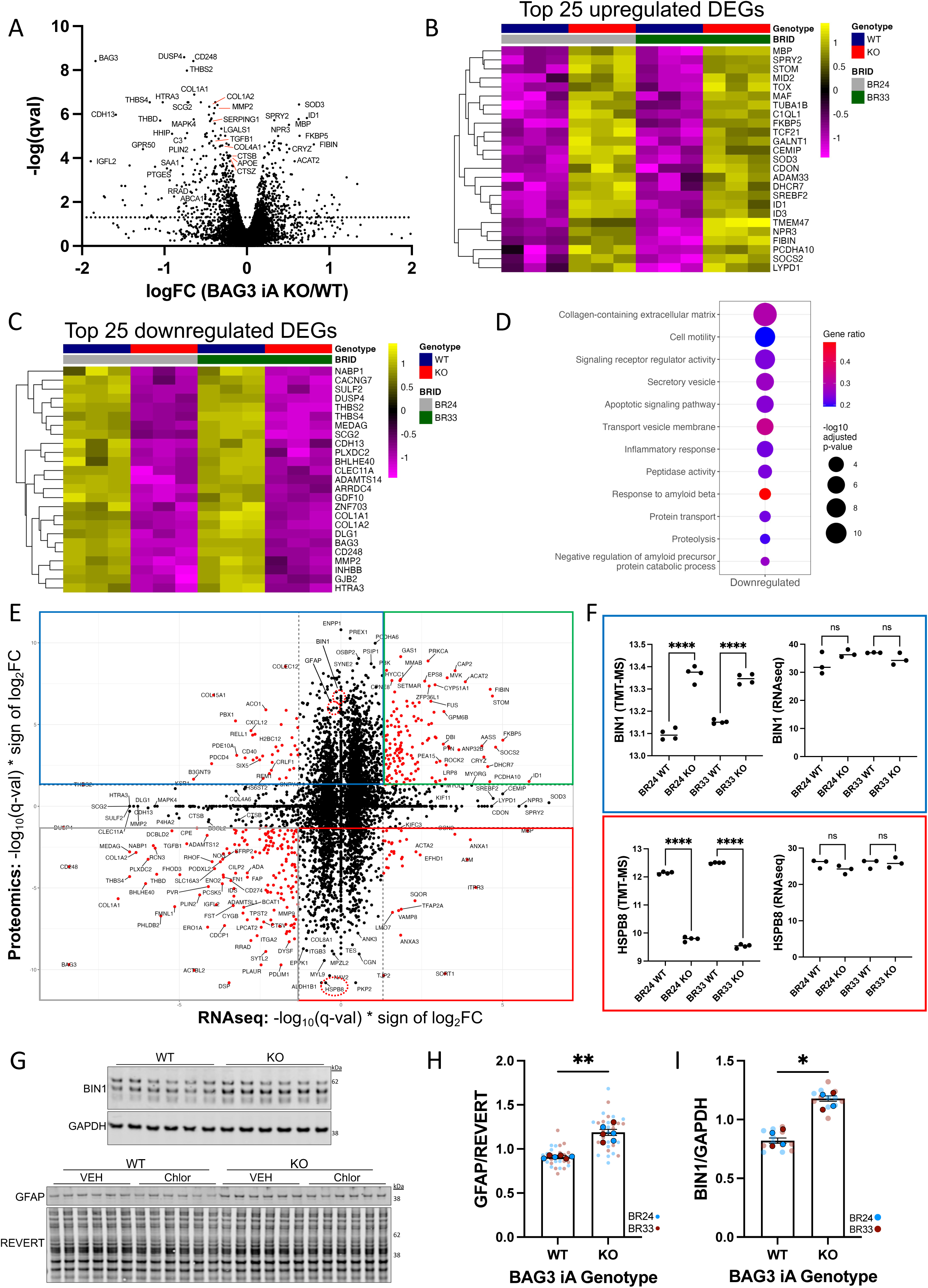
Integration of transcriptomics and proteomics following BAG3 loss reveals post-transcriptional control of genes and pathways relevant to neurodegeneration. (**A**) Volcano plot of BAG3 WT vs KO iAs, showing differentially expressed genes (DEGs) as measured by RNAseq (dots above the dotted line; q < 0.05). Data shown includes BR24 and BR33 combined, with line ID regressed away (n = 4 wells per genetic background). (**B**) Heat map showing the top 25 upregulated DEGs in BAG3 KO iAs across both genetic backgrounds following RNAseq analysis. (**C**) Heat map showing the top 25 downregulated DEGs in BAG3 KO iAs across both genetic backgrounds following RNAseq analysis. (**D**) Gene Ontology gene set enrichment analysis showing the downregulated pathways in BAG3 KO iAs. Derived from the RNAseq. (**E**) Integration plot of the differentially expressed proteins in Figure 1F and the DEGs identified from RNAseq. Red dots represent significance (q < .05) at both the protein and RNA level. Genes of interest are encircled with a red dotted line. (**F**) Representative graphs of BIN1 (protein up, RNA unchanged) and HSPB8 (protein down, RNA unchanged), genes found in the segmented quadrants in Figure 5E where RNA and protein are discordant. Data derived from the TMT-MS in Figure 1F or RNAseq. Statistics for the RNA and protein abundance were performed using an unpaired one-way ANOVA with Tukey’s post hoc testing. (**G**) Representative WBs of BIN1 normalized to GAPDH and GFAP normalized to REVERT in BAG3 WT and KO astrocytes. (**H**) Quantification of GFAP normalized to REVERT in BAG3 WT and KO astrocytes (n = 6 differentiations across 2 genetic backgrounds and 4 technical replicates per differentiation; paired t-test). (**I**) Quantification of BIN1 normalized to GAPDH in BAG3 WT and KO astrocytes (n = 4 differentiations across 2 genetic backgrounds and 2-4 technical replicates per differentiation; paired t-test). Data is displayed using SEM. Each solid dot corresponds to a single differentiation with technical replicates (transparent smaller dots) averaged. Blue = BR24, Red = BR33. For all comparisons: \**p* < 0.05, \*\**p* < 0.01, \*\*\**p* < 0.001, \*\*\*\**p* < 0.0001, ns = not significant.

Genes exhibiting discordance between RNA and protein expression were of particular interest to the role of BAG3 in proteostasis because such discrepancies suggest BAG3 may exert its regulatory influence post-transcriptionally, primarily at the protein level (**Fig. 5E-F**). Considering the observed alterations in protein degradation and the interacting proteins related to degradation and the retromer, we focused our validation on the proteins in these quadrants.

Two of the proteins chosen for validation via WB analyses were BIN1, an AD GWAS gene, and GFAP, a protein important for astrocyte reactivity and AD. Indeed, both BIN1 and GFAP were validated via WB and shown to be increased at the protein level following BAG3 KO (**Figure 5G-I, Supplementary Fig. S7A-B**).

### BAG3 regulates HSPB8 protein levels and ubiquitinated protein clearance under proteasome stress

BAG3 was previously shown to physically bind to HSPB8 in multiple cell types [8], and we detected this binding also within astrocytes. While HSPB8 levels were fully ablated in BAG3 KO astrocytes, this has not been reported previously, indicating this regulatory/destabilizing effect could be of particular relevance in astrocytes (**Fig. 5F**). To determine whether HSPB8 could be restored by reintroducing BAG3, we overexpressed BAG3 in both WT and KO astrocytes using either BAG3 WT or an empty vector control. While HSPB8 levels were partially rescued in KO astrocytes, the effect was modest, likely due to limited BAG3 overexpression efficiency which only rescued ∼25% of endogenous BAG3 levels (**Fig. 6A-C, Supplementary Fig. S7C-D**). We then impaired proteasome and autophagy-mediated protein degradation pathways in BAG3 WT and KO astrocytes by treating with bortezomib or chloroquine, respectively, and measured HSPB8 abundance by WB. Chloroquine did not alter HSPB8 protein levels, whereas bortezomib significantly increased HSPB8 abundance in both WT and KO astrocytes (**Fig. 6D-E, Supplementary Fig. S7E**), suggesting this accumulation occurs independently of BAG3 and may be a direct result of proteasome inhibition. Given this observation, we took a more agnostic approach to assess whether global ubiquitinated protein turnover was altered in WT versus KO astrocytes. While no differences were observed at baseline, bortezomib treatment led to a greater accumulation of ubiquitinated protein in KO compared to WT astrocytes, indicating that BAG3 loss broadly impairs protein clearance during proteasome stress (**Fig. 6F-G, Supplementary Fig. S7F**).

**Figure 6.**
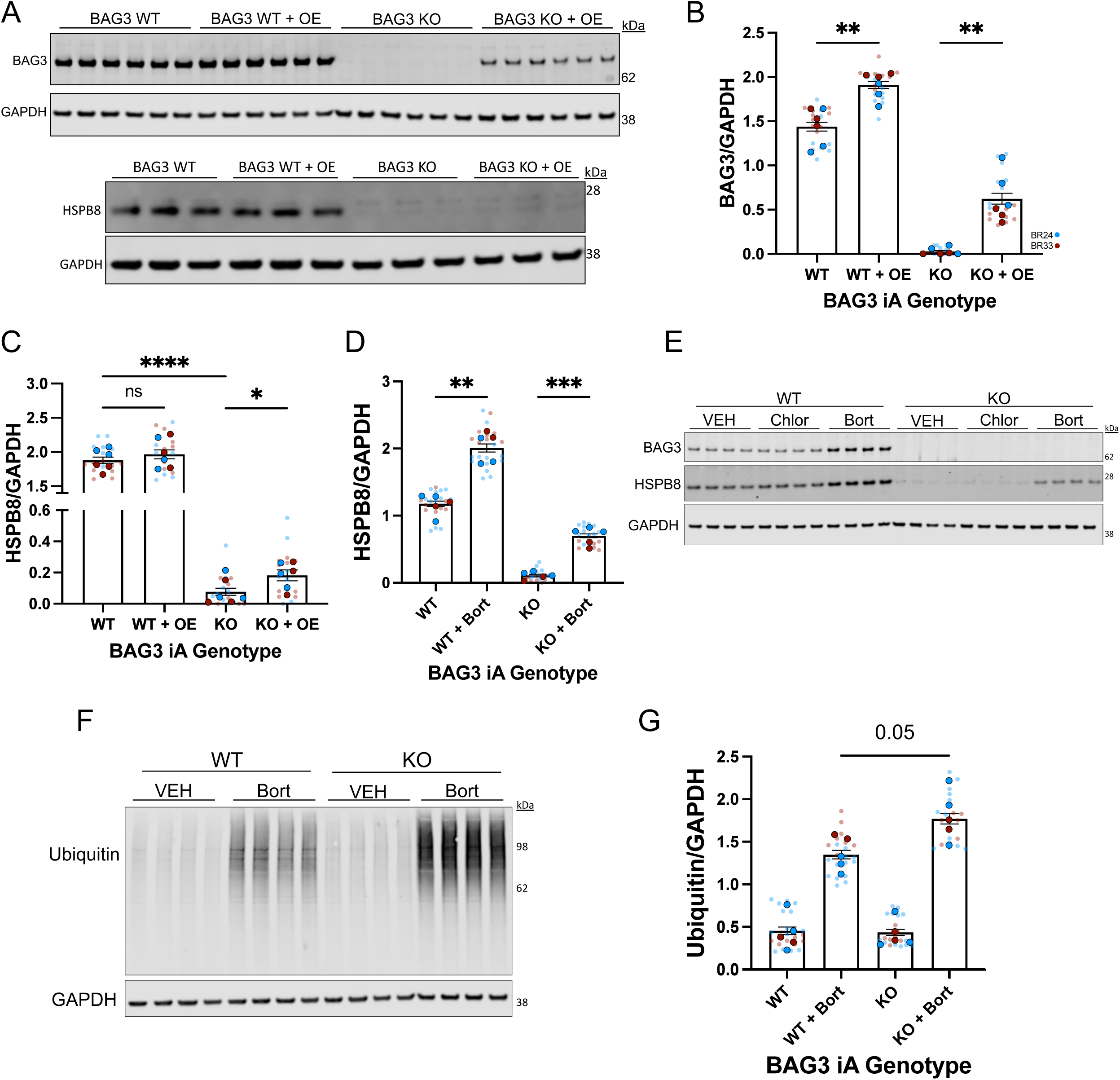
BAG3 regulates HSPB8 and ubiquitinated protein clearance under proteasome stress. (**A**) Representative western blots (WB) showing HSPB8 and BAG3 protein level changes in BAG3 WT and KO iAs as well as BAG3 overexpression (OE) iAs. (**B**) Quantification of BAG3 normalized to GAPDH following KO and OE (n = 6 differentiations across 2 genetic backgrounds and 3 technical replicates per differentiation). (**C**) Quantification of HSPB8 normalized to GAPDH following KO and OE of BAG3 (n = 6 differentiations across 2 genetic backgrounds and 3 technical replicates per differentiation). (**D**) HSPB8 protein abundance following bortezomib treatment (5 nM for 24 hours) in BAG3 WT and KO iAs (n = 5 differentiations across 2 genetic backgrounds and 3 technical replicates per differentiation). (**E**) Representative WB showing BAG3 and HSPB8 following bortezomib (5 nM for 24 hours) or chloroquine (80 µM for 24 hours) treatment. (**F**) Representative WB showing total ubiquitinated protein in BAG3 WT and KO iAs with our without bortezomib treatment (5 nM for 24 hours). (**G**) Total ubiquitinated protein was quantified and normalized to GAPDH (n = 5 differentiations across 2 genetic backgrounds, and 3-6 technical replicates per differentiation). For all graphs, data is displayed using SEM and for statistical analysis, a paired one-way ANOVA with Tukey’s post hoc testing was used. Each solid dot corresponds to a single differentiation with technical replicates (transparent smaller dots) averaged. Blue = BR24, Red = BR33. For all comparisons: \**p* < 0.05, \*\**p* < 0.01, \*\*\**p* < 0.001, \*\*\*\**p* < 0.0001, ns = not significant.

### Relevance of BAG3 in astrocytes to Alzheimer’s disease: BAG3 defines a specific stress-response subtype of astrocytes within the human brain and loss of BAG3 reduces amyloid-**β** 42 clearance

Considering the relevance of BAG3 in astrocytic protein degradation and functioning as well as the observation that BAG3 KO reduces the transcripts of genes related to amyloid beta response, we next determined how these phenotypes may coalesce to affect AD aggregation prone proteins. We began by treating astrocytes with aqueous-soluble AD brain extract. This extract preparation has been used in numerous studies as an AD-relevant stressor [4, 20, 21, 40, 48]. Across iAs from nine individuals in the ROSMAP cohorts of aging, BAG3 protein abundance significantly increased in response to AD brain extract exposure, as measured by WB (**Fig. 7A**). After confirming BAG3 responds to AD-relevant stimuli in iAs, we examined astrocyte specific BAG3 effects on fAD neurons. We had previously observed that fAD mutant iNs express higher levels of BAG3 protein compared to their isogenic controls, consistent with the upregulation of BAG3 observed in iAs with exposure to AD brain extract (Augur *et. al.*, in revision). Here, we determined whether astrocytic BAG3 abundance affects Aβ clearance when cocultured with APP/PSEN1 mutant iNs. To test this, we established cocultures of APP/PSEN1 mutant iNs with either BAG3 WT or KO iAs for 6 days and measured Aβ peptides in the culture medium (**Fig. 7B-C**). Strikingly, Aβ42 levels, but not Aβ38 or Aβ40 were significantly elevated in the conditioned medium of co-cultures containing BAG3 KO iAs, indicating impaired clearance of Aβ42 (**Fig. 7D-F**).

**Figure 7.**
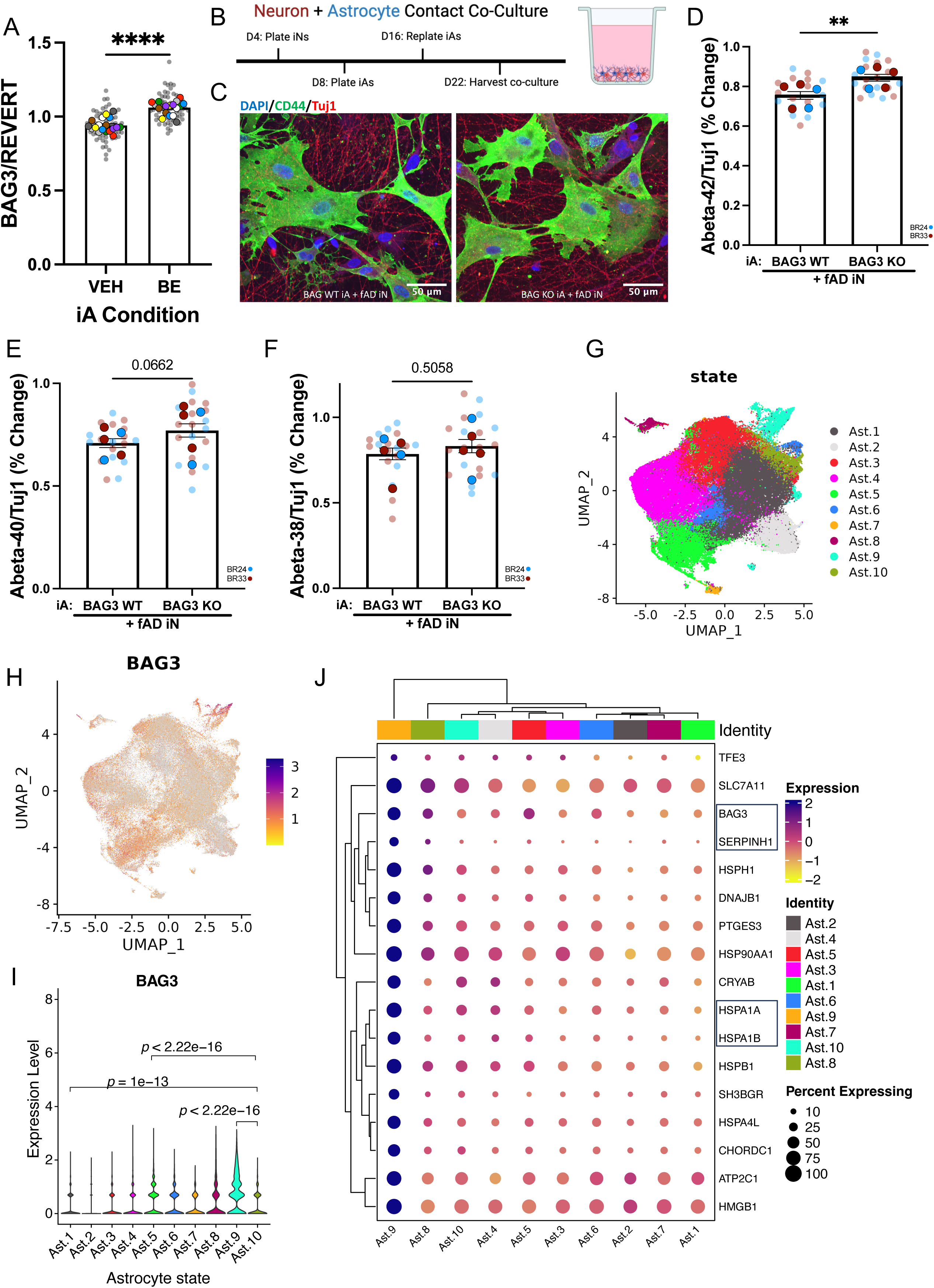
Loss of BAG3 in astrocytes reduces amyloid-β 42 clearance from APP/PSEN1 mutant neurons and BAG3 defines a specific subtype of astrocytes within the human brain. (**A**) iAs were treated for 72 hours with AD human brain extract in cell culture media. The upregulation of BAG3 following exposure to brain extract was quantified via western blot (WB). BAG3 was normalized to total protein using REVERT. Solid colored dots represent a differentiation while small gray dots in the background represent all technical replicates (n = 18 differentiations across 9 genetic backgrounds and 3-4 technical replicates per differentiation). (**B**) Visual timeline for iN and iA contact cocultures showing 6-day cocultures beginning on day 16 of iN and iA differentiation and continuing until day 22. (**C**) Representative fluorescent images of iN and iA 6-day cocultures. fAD iNs were cocultured with BAG3 WT or BAG3 KO iAs. CD44 antibody was used to show astrocyte morphology while Tuj1 staining depicts neurons. Scale bar = 50 µm. (**D**) Aβ 42, (**E**) Aβ 40, and (**F**) Aβ 38 were measured in the 48 hour condition media of contact cocultures via an MSD Aβ ELISA. The percent change of extracellular Aβ in the condition media compared to fAD iN monocoluture was normalized to neuron abundance using the neuron marker, Tuj1 (n = 5 differentiations across 2 genetic backgrounds and 3-4 technical replicates per differentiation). (**G**) Uniform Manifold Approximation Projection (UMAP) embedding of astrocytes calculated based on snRNAseq data derived from 437 ROSMAP dorsolateral prefrontal cortex samples. These astrocytes were previously partitioned into 10 transcriptionally distinct subclusters (Ast.1-Ast.10) [15]. (**H**) The same UMAP is shown as in G, except each cell is colored by their log-normalized BAG3 expression. (**I**) BAG3 transcript expression (log-normalized) compared across the 10 astrocyte subclusters (p-values: two-sided unpaired t-test) (**J**) Top astrocyte subcluster 9 (Ast.9) marker gene expression compared to all other astrocyte subclusters. (Color bar: scaled gene expression; Circle size: percent of cells expressed). Where relevant, data is displayed using SEM and for statistical analysis, a paired t-test was used. Each solid dot corresponds to a single differentiation with technical replicates (transparent smaller dots) averaged. Blue = BR24, Red = BR33. For all comparisons: \**p* < 0.05, \*\**p* < 0.01, \*\*\**p* < 0.001, \*\*\*\**p* < 0.0001, ns = not significant.

To extend our findings to the human brain, we again probed the snRNA-seq dataset utilized in Figure 1. As described previously [15], astrocytes in this snRNA-seq dataset were partitioned into 10 transcriptionally distinct subpopulations. One of these subpopulations, Astrocyte Subcluster 9 (Ast.9), was enriched for stress-responsive genes, including multiple heat shock proteins known to interact with BAG3 and that appear in our BAG3 interactome analysis (**Fig. 7G–I**). BAG3 was among the top defining markers for Ast.9 and was uniquely upregulated in Ast.9 compared to all other astrocyte subclusters (adjusted p-value range: 10^-79^ – 2.0 x 10^-3^) [12], further supporting its specialized role in the astrocytic stress response state in AD and the human brain (**Figure 7J**). Taken together, these observations suggest that BAG3 is poised to impact proteostasis, Aβ clearance, and cognitive outcomes in the aged human brain within a subtype of astrocytes (**Supplementary Fig. S8**).

## DISCUSSION

BAG3 is emerging as a critical player in cellular proteostasis and selective autophagy, yet its precise role in astrocyte biology and neurodegeneration has remained poorly defined. In this study, we provide evidence that BAG3 is not only enriched in human astrocytes but also modulates essential aspects of their proteostatic machinery. Using a combination of CRISPR-edited iPSC-derived models, transcriptomic and proteomic profiling, mechanistic validation, and postmortem human brain tissue analysis, we present a model in which BAG3 supports astrocyte function through interactions with key binding partners, facilitation of autophagic-lysosomal degradation, modulation of ubiquitin-proteasome system (UPS) activity, and regulation of ubiquitinated protein substrates (**Fig. 8**).

**Figure 8.**
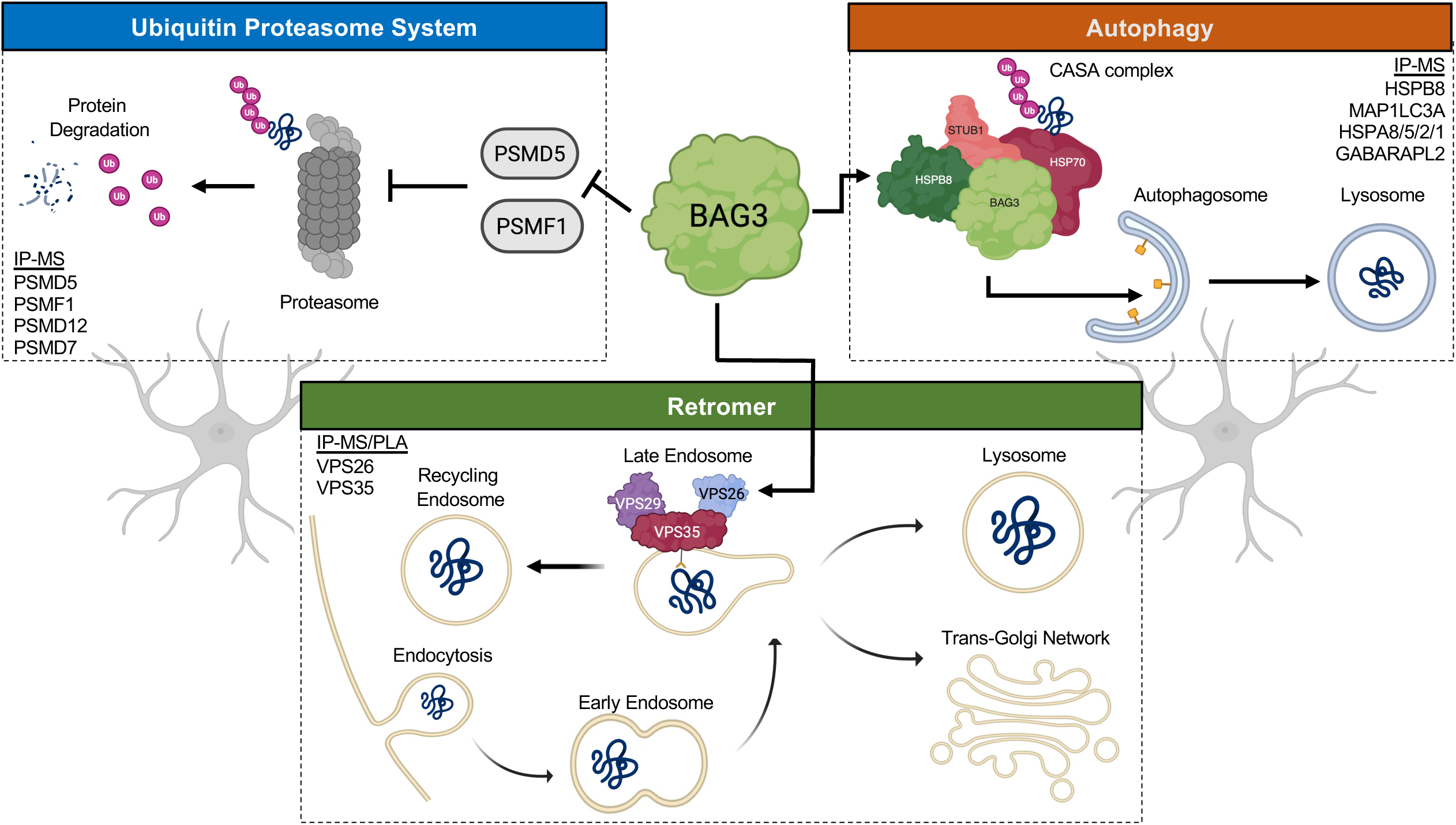
BAG3 is a molecular hub at the center of the protein quality control network in astrocytes. A representative image showing how BAG3 interacts with the ubiquitin proteasome system (UPS), autophagy, and the retromer. We hypothesize BAG3 directly interacts with and sequesters proteasome inhibitors (PSMD5, PSMF1, etc.) to modulate UPS activity, interacts with the retromer complex via VPS26 or VPS35 binding, and significantly alters autophagy and lysosomal function and numbers within astrocytes.

A striking finding of this work is the extent of proteomic disruption following BAG3 ablation in astrocytes. Compared to neurons, BAG3 knockout (KO) in astrocytes resulted in significantly more differentially expressed proteins (DEPs), suggesting a unique dependence on BAG3-mediated proteostasis in these glial cells. This vulnerability may stem from their higher BAG3 and HSPB8 expression and its central role in extracellular protein clearance. BAG3 KO astrocytes exhibited marked depletion of autophagy-lysosomal pathway proteins, reduced lysosome numbers, and impaired degradation capacity, features consistent with compromised proteostatic control. Comparing astrocyte reliance on other autophagy pathways to the pronounced effects of BAG3 knockout would be an interesting future direction, as these results might further elucidate the importance of BAG3 mediated autophagy in these cells.

While the role of BAG3 in the chaperone-assisted selective autophagy (CASA) complex is well established, we found a cell-type specific dependency of HSPB8 on BAG3. BAG3 deficiency led to a post-transcriptional loss of HSPB8 protein in astrocytes, a phenomenon not previously reported in other cell types to our knowledge. This supports BAG3’s role as a scaffold for the CASA complex and further positions it as a key mediator of autophagy in astrocytes.

Reintroduction of lower levels of BAG3 (∼25% of WT) only partially restored HSPB8 levels. However, the HSPB8 response to proteasome inhibition with bortezomib was preserved, indicating that the heat shock response remains at least partially intact in BAG3-deficient astrocytes. It also cannot be excluded that proteasomal degradation may contribute to HSPB8 loss in this context. This finding is particularly significant given the diverse cellular roles of HSPB8, which evidently may depend on sufficient BAG3 functioning in astrocytes.

Beyond HSPB8, we identified broader post-translational regulatory roles for BAG3, which could influence AD and other neurodegenerative diseases. Notably, BIN1 protein levels were elevated in BAG3 KO astrocytes without corresponding transcript changes, pointing to a role for BAG3 in limiting accumulation of specific membrane trafficking proteins. Given the involvement of BIN1 in endocytosis, tau propagation, and its status as an AD GWAS gene, its dysregulation in BAG3-deficient astrocytes could contribute to AD pathology. Similarly, GFAP was upregulated following BAG3 loss, suggesting that BAG3 may have a role in modulating astrocyte reactivity, as has been suggested previously [44]. At the transcript level, aggregation-prone proteins like FUS were altered in BAG3 KO astrocytes, implicating BAG3 in broader transcriptional level alterations. These phenotypes highlight multiple BAG3-regulated pathways worthy of future investigation.

One of the most important implications of this study is that it supports the hypothesis that BAG3 serves as a molecular hub at the intersection of autophagy, the UPS, and the retromer complex (**Fig. 8**). This suggests BAG3 is a critical node within the protein quality control and intracellular trafficking network of astrocytes. We show a novel interaction between BAG3 and the core retromer complex, supported by IP-MS, AlphaFold structural modeling, and PLA. Specifically, we identify a predicted interaction between the IPV domain of BAG3 (centered on isoleucine 208) and a hydrophobic pocket within VPS26. Notably, this same IPV domain mediates BAG3 binding to small heat shock proteins like HSPB8. One way to interpret this data is to propose a competitive binding model in which BAG3 acts as a molecular switch. Under basal conditions, it could preferentially bind the retromer to support receptor recycling and retrograde transport.

Under proteotoxic stress, HSPB8 may outcompete the retromer for BAG3 binding, thereby shifting astrocytic machinery toward autophagic degradation. This switch-like mechanism could allow astrocytes to dynamically prioritize intracellular trafficking/recycling versus degradation depending on proteostatic demands. Although our IP-MS and modeling data do not fully resolve the interaction dynamics with the retromer complex, they provide compelling initial support for BAG3’s involvement in this pathway. This model is bolstered by growing literature implicating both retromer dysfunction and BAG3 dysregulation in the pathogenesis of Alzheimer’s disease, Parkinson’s disease, and frontotemporal dementia, disorders characterized by impaired protein clearance and trafficking [50]. Astrocytes, with their roles in neuronal support, metabolism, and extracellular aggregate clearance, may be especially vulnerable to BAG3-mediated dysfunction. Intriguingly, BAG3 and VPS35 have both emerged as risk genes in PD GWAS studies, suggesting potential convergence on a causal mechanism [16, 32].

Extending our hypothesis that BAG3 functions as a central regulator of proteostasis, we uncover, for the first time, a direct role for BAG3 in modulating the UPS in astrocytes. Specifically, we validate BAG3 interactions with two key UPS regulatory proteins, PSMF1 and PSMD5, through IP-MS. These interactions appear to be specific to regulatory components of the proteasome, as no interaction was observed between BAG3 and the core proteasome component, PSMB5. This suggests that BAG3 may influence UPS activity through indirect regulatory mechanisms rather than direct engagement with the proteasome catalytic core. One hypothesis is that BAG3 may temper the effect of PSMD5 on proteosome assembly during basal and stress conditions, which would maintain efficient proteosome activity [27, 42].

Similarly, PSMF1 has been described to have inhibitory effects on the proteosome, and BAG3 binding may inhibit this function [11, 46]. Further supporting the functional significance of these interactions, we found that expression of the disease-associated BAG3 E455K mutant abolished the interaction between BAG3 and PSMD5. This point mutation, previously identified in patients with cardiomyopathy and neuropathy, may therefore impair BAG3-mediated regulation of the UPS, an effect with potential clinical relevance that warrants deeper investigation. Functionally, BAG3 knockout in astrocytes led to a marked reduction in UPS chymotrypsin-like activity, indicating compromised proteasome function, which would support the role of BAG3 regulating UPS inhibitory proteins. Conversely, BAG3 overexpression in HEK293FT cells enhanced UPS activity, reinforcing the conclusion that BAG3 promotes proteasomal degradation capacity, potentially via its sequestration of PSMF1 or PSMD5.

Finally, we demonstrate that BAG3 loss in astrocytes decreases genes related to the transcriptomic response to Aβ, as seen in our RNAseq profiling, as well as the clearance of Aβ42 in co-culture systems with fAD neurons. In the context of APP/PSEN1 driven Aβ overproduction, BAG3-deficient astrocytes exhibit a reduction in their capacity to clear extracellular Aβ42, and to a lesser extent Aβ40, highlighting a previously underappreciated role for astrocytic BAG3 in Aβ proteostasis. While previous studies have predominantly focused on the role of BAG3 in neuronal tau clearance, relatively few have examined the influence of BAG3 on Aβ clearance. The exact mechanism underlying altered Aβ clearance requires further investigation, as only limited changes were detected using the endocytosis assays employed. Alterations in other forms of receptor-mediated or clathrin-mediated endocytosis cannot be ruled out with existing data.

Our findings provide novel evidence that BAG3-mediated proteostatic mechanisms in astrocytes are essential for maintaining efficient Aβ clearance and suggest that BAG3 upregulation in astrocytes may be a direct result of AD pathologies in the brain. Supporting this idea, snRNA-seq analysis from postmortem human brain tissue from over 400 individuals reveals that BAG3 is selectively enriched in a reactive astrocyte subpopulation (Ast.9), which also expresses several heat shock proteins and stress-responsive chaperones. We hypothesize that this astrocyte subtype may colocalize with sites of pathological protein accumulation, potentially representing a BAG3-driven stress response state triggered by chronic proteotoxic burden.

## CONCLUSION

Taken together, these data converge on a model in which BAG3 acts as a central regulator of astrocytic adaptation to neurodegenerative stress like Aβ. The integration of iPSC-based functional assays, quantitative proteomics, interactome analyses, and human single-cell transcriptomics support the hypothesis that BAG3 could serve as a molecular hub orchestrating astrocytic response to chronic proteinopathy in Alzheimer’s disease. Failure to efficiently upregulate BAG3 may impair the brain’s ability to clear unwanted proteins and contribute to neurodegeneration. The data presented herein identifies specific proteins and pathways that mediate the effects of BAG3, which represent candidate targets for therapeutic intervention and provide a foundation for future work in this area.

## DECLARATIONS

### Ethics approval and consent to participate

Informed consent was obtained by all human participants with proper IRB approvals.

### Consent for publication

Not applicable.

### Availability of data and material

The datasets supporting the conclusions of this article are included within the article (and its supplemental files).

### Authors’ contributions

Z.M.A and T.L.Y.-P. conceptualized the project design and methodology. Z.M.A. conducted the bulk of experimentation and analyses and wrote the manuscript. G.M.F., Z.R.M., G.T., and M.R.A., contributed to experimentation and analyses in the manuscript. C.R.B. assisted in data curation and computational analyses. N.C.-L. and P.L.D-J. assisted in snRNAseq analyses and data interpretation. D.M.D and N.T.S. contributed to the proteomic data acquisition and analysis.

## LIST OF ABBREVIATIONS

BAG3: Bcl-2-associated athanogene 3
AD: Alzheimer’s disease
pTau: phosphorylated tau
ALP: autophagy-lysosomal pathway
iPSC: induced pluripotent stem cell
iNs: iPSC-derived neurons
iAs: iPSC-derived astrocytes
KO: knockout
WT: wildtype
Aß: amyloid-ß
ROS and MAP: Religious Orders Study and Rush Memory and Aging Projects
TBS: tris-buffered saline
SDS: sodium dodecyl sulfate
WB: Western blot
GSEA: gene set enrichment analysis
DEPs: differentially expressed proteins
DEGs: differentially expressed genes
DIA-MS: data-independent acquisition mass spectrometry
TMT-MS: tandem mass tag mass spectrometry
GO: gene ontology
IP-MS: immunoprecipitation mass spectrometry
snRNAseq: single nucleus RNA sequencing
ICC: immunocytochemistry
PAE: predicted aligned error
PLA: proximity ligation assay
VEH: vehicle
Bort: bortezomib
Chlor: chloroquine
H.S.: heat shock

## Acknowledgements

The authors would like to thank Richard Zhu for his computational support. We also thank the NeuroTechnology Studio at Brigham and Women’s Hospital for providing Zeiss LSM710, Andor Dragonfly 600 Spinning Disk, BD LSRFortessa^TM^ Cell Analyzer, and Incucyte S3 Live-Cell Analysis System instrument access and consultation on data acquisition and analysis; the iPSC NeuroHub at Brigham and Women’s Hospital for technical assistance; and the Proteomics Core at Emory University for proteomic analysis and consultation. Z.M.A. thanks Stephen Haggarty, Mel Feany, and Daniel Finley for their Dissertation Advisory Committee guidance and to all members of the Tracy Young-Pearse lab, we thank them for their invaluable feedback and support throughout the duration of this project.

## SUPPLEMENTARY FIGURE LEGENDS

**Supplementary Figure 1.** Generation of BAG3 isogenic lines using CRISPR/Cas9. (**A**) Graphic showing the domain architecture of BAG3 protein as well as the guides used for CRISPR/Cas9 editing. (**B**) Table listing the BAG3 isogenic lines generated in this study. *BAG3* WT and KO iPSCs were generated in two isogenic sets of lines: BR33 (male) and BR24 (female) derived from non-cognitively impaired participants in the ROSMAP cohort. CRISPR/Cas9 editing was performed at the iPSC stage with a gRNA targeted to exon 2 of BAG3, followed by monoclonal selection and sequencing. For analysis, two clones for each iPSC line were selected: one targeted (biallelic frameshift mutations, KO) and an unaltered wildtype clone (WT). (**C**) Full, uncropped representative western blot of Figure 1B.

**Supplementary Figure 2.** BAG3 pathway analysis in ROSMAP iAs and iNs. (**A**) Gene ontology pathway analysis using genes identified following (**B**) gene (RNAseq) and protein (TMT-MS) level Pearson correlations with *BAG3.* Each dot represents data for a single gene (n = 43 ROSMAP iA lines) [25]. Left, genes positively correlated (> +0.4 Pearson r) to BAG3 at both the protein and RNA level across ROSMAP iAs (upper right quadrant). Right, genes negatively correlated (< –0.4 Pearson r) to BAG3 at both the protein and RNA level across ROSMAP iAs (lower left quadrant). (**C**) Gene (RNAseq) and protein (TMT-MS) level Pearson correlations with neuronal *BAG3* were plotted and each dot represents data for a single gene (n = 53 ROSMAP iN lines) [19]. (**D**) Gene Ontology (GO) pathway analysis using genes identified in (C). Left, genes positively correlated (> +0.4 Pearson r) to BAG3 at both the protein and RNA level across ROSMAP iNs (upper right quadrant). Right, genes negatively correlated (< –0.4 Pearson r) to BAG3 at both the protein and RNA level across ROSMAP iNs (lower left quadrant). (**E**) GO pathway enrichment following gene partitioning across BAG3 WT vs KO iAs (q < 0.05, log_2_FC > –0.2). For correlation with BAG3 in ROSMAP iAs there must be a positive BAG3 RNA and protein correlation across 43 ROSMAP iA proteomes and transcriptomes (Pearson r > 0.4 for both RNA and protein). For correlation in ROSMAP brains, there must be a positive BAG3 protein correlation across 971 ROSMAP human brains (q-value < 0.05, Pearson r > 0.2). (**F**) Gene concept network of genes downregulated in BAG3 KO iAs that belong to the actin filament binding, phospholipid binding, and peptidase regulator activity pathways shown in (E).

**Supplementary Figure 3.** Predicted aligned error (PAE) plots of BAG3 binding partners in astrocytes. PAE plots of BAG3 and several proteins of interest identified in the iA coIP-MS. PAE plots were created using AlphaFold Multimer and Model 1 of 5 is shown with consistency observed across all models. The proteins shown include (**A**) VPS29 and (**B**) VPS35. (**C**) Representative fluorescent images of a proximity ligation assay between the empty vector control and VPS26. Red puncta would represent positive signal or proximity between the two proteins. GFP positive cells represent positive plasmid transduction. (**D**) PAE plot of BAG3 and PSMB5. (**E-G**) Full, uncropped representative western blots showing the co-immunoprecipitation of proteins of interest from Figure 2B-D.

**Supplementary Figure 4.** Uncropped representative western blots of HEK293FT BAG3 co-immunoprecipitation. Full, uncropped representative western blots showing the co-immunoprecipitation of (**A**) BAG3, (**B**) PSMD5, (**C**) VPS35, (**D**) PSMF1, and (**E**) PSMB5 from Figure 3A.

**Supplementary Figure 5.** Endocytosis phenotypes are inconsistent with loss of BAG3. (**A-B**) RAB5 puncta per astrocyte cell area, as defined by CD44 staining, were quantified using cell-profiler (n = 2 differentiations across 2 genetic backgrounds and 3 technical replicates per differentiation with 8 images taken per replicate). (**C-D**) pHrodo-Dextran measured across BAG3 WT and KO iAs with flow cytometry. Average signal per cell was calculated for each genetic background and an untreated control (n = 6 differentiations across 2 genetic background). (**E**) BSA-GFP measured across BAG3 WT and KO iAs with flow cytometry. Average signal per cell was calculated for each genetic background (n = 2 differentiations across 2 genetic backgrounds with 2 technical replicates each). For all comparisons: \**p* < 0.05, \*\**p* < 0.01, \*\*\**p* < 0.001, \*\*\*\**p* < 0.0001, ns = not significant. Full, uncropped representative western blots of (**F**) Figure 4B and (**G**) Figure 4I.

**Supplementary Figure 6.** Transcriptional level changes in BAG3 KO astrocytes. (**A**) Principal component analysis (PCA) plot of the BAG3 WT vs KO iA RNAseq following line ID regression and normalization. (**B**) Gene set enrichment analysis showing several of the most interesting, upregulated pathways from the RNAseq comparing BAG3 WT and KO iAs. (**C**) Representative graphs of FUS (protein up, RNA up) and ACTBL2 (protein down, RNA down), genes found in the segmented quadrants in Figure 5E where RNA and protein are concordant. Statistics for the RNA and protein abundance were performed using an unpaired one-way ANOVA with Tukey’s post hoc testing. (**D**) Representative western blot images of HSF1 transcription factor in BAG3 WT and KO iAs following heat shock (H.S.) or 5 nM Bortezomib (Bort) treatment. (**E-F**) HSF1 positive astrocyte nuclei were calculated with ICC imaging. The percent of blue nuclei with overlaying red HSF1 staining was calculated for BAG3 WT and KO astrocytes (n = 4 differentiations across 2 genetic backgrounds with 6-8 technical replicates per differentiation; SEM with a paired t-test). Each solid dot corresponds to a single differentiation with technical replicates (transparent smaller dots) averaged. Blue = BR24, Red = BR33. For all comparisons: \**p* < 0.05, \*\**p* < 0.01, \*\*\**p* < 0.001, \*\*\*\**p* < 0.0001, ns = not significant.

**Supplementary Figure 7.** Uncropped representative western blots. Full, uncropped representative western blots of (**A**) BIN1 in Figure 5G, (**B**) GFAP in Figure 5G, (**C-D**) BAG3 OE from Figure 6A, (**E**) BAG3 and HSPB8 with Bort and Chlor treatment in Figure 6E, (**F**) ubiquitin following Bort treatment in Figure 6F.

**Supplementary Figure 8.** Ast.9 subcluster abundance correlates with cognitive measures. (**A**) MMSE, (**B**) Cognition, (**C**) Amyloid, (**D**) and Tangles scoring was determined for the 20 individuals in the ROSMAP cohort of individuals that had Ast.9 abundance in the brain. Pearson correlations were performed for each comparison with the r and p values shown for each.

