## Supplemental Figures for "BAG3 coordinates astrocytic proteostasis of Alzheimer’s disease-linked proteins via proteasome, autophagy, and retromer complex interactions"

### Supplementary Fig. S1

A

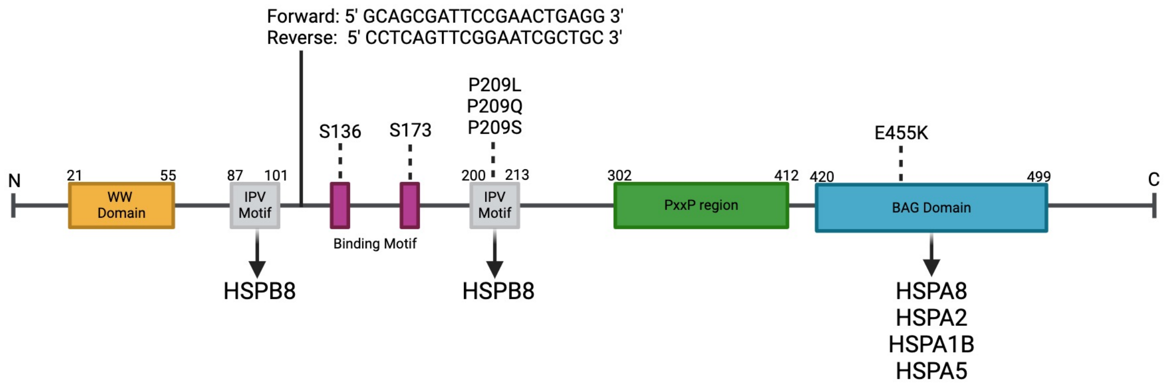


B

| Line ID | Clone | Allele | Editing | Genotype |
|---|---|---|---|---|
| BR24<br>Age: 90<br>Sex: Female<br>APOE: 3/3 | 16 | 1 | N/A | WT |
| | | 2 | N/A | |
| | 14 | 1 | 2bp insertion | KO |
| | | 2 | 2bp insertion | |
| BR33<br>Age: 91<br>Sex: Male<br>APOE: 3/3 | 12 | 1 | N/A | WT |
| | | 2 | N/A | |
| | 21 | 1 | 2bp insertion | KO |
| | | 2 | 2bp insertion | |

C

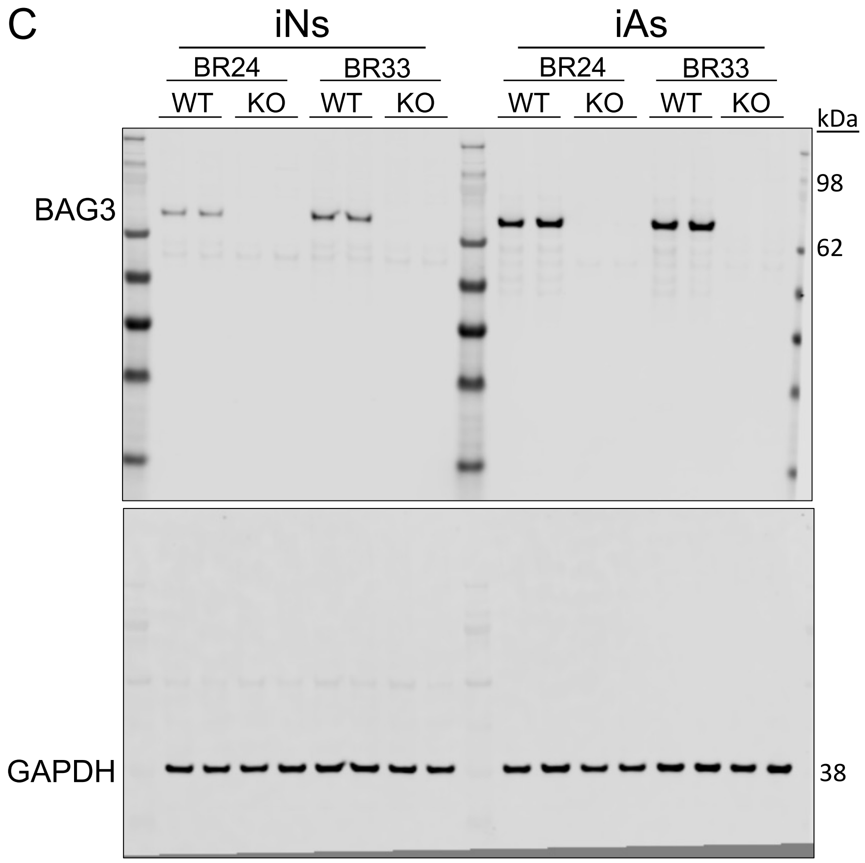

### Supplementary Fig. S2

A

#### ROSMAP iAs

| Term | Count | % | p-value | Adj. p-value |
|---|---|---|---|---|
| Protein Transport | 56 | 12.1 | $6.3 \times 10^{-27}$ | $1.4 \times 10^{-23}$ |
| ER to Golgi Vesicle-Mediated Transport | 30 | 6.5 | $5.8 \times 10^{-21}$ | $5.7 \times 10^{-18}$ |
| Intracellular Protein Transport | 41 | 8.8 | $7.5 \times 10^{-21}$ | $5.7 \times 10^{-18}$ |
| Endocytic Recycling | 18 | 3.9 | $2.1 \times 10^{-13}$ | $1.2 \times 10^{-10}$ |
| COPII-Coated Vesicle Cargo Loading | 10 | 2.2 | $2.6 \times 10^{-11}$ | $1.2 \times 10^{-8}$ |
| Vesicle-Mediated Transport | 24 | 5.2 | $1.7 \times 10^{-9}$ | $5.4 \times 10^{-7}$ |
| Actin Cytoskeleton Organization | 22 | 4.7 | $3.9 \times 10^{-8}$ | $8.7 \times 10^{-6}$ |
| Golgi Organization | 17 | 3.7 | $4.2 \times 10^{-8}$ | $8.7 \times 10^{-6}$ |
| Endocytosis | 17 | 3.7 | $1.7 \times 10^{-6}$ | $3.3 \times 10^{-4}$ |
| Retrograde Transport | 5 | 1.1 | $4.6 \times 10^{-5}$ | $7.5 \times 10^{-3}$ |

Positive Correlation (+0.4 Pearson or higher)

| Term | Count | % | p-value | Adj. p-value |
|---|---|---|---|---|
| rRNA Processing | 87 | 6.8 | $4.3 \times 10^{-73}$ | $1.3 \times 10^{-69}$ |
| Translation | 106 | 8.3 | $4.1 \times 10^{-72}$ | $6.0 \times 10^{-69}$ |
| mRNA Splicing | 104 | 8.1 | $1.2 \times 10^{-69}$ | $9.0 \times 10^{-67}$ |
| Ribosomal Small Subunit Biogenesis | 66 | 5.1 | $2.8 \times 10^{-67}$ | $1.6 \times 10^{-64}$ |
| DNA Repair | 68 | 5.3 | $1.9 \times 10^{-23}$ | $6.9 \times 10^{-21}$ |
| Cell Division | 83 | 6.5 | $2.3 \times 10^{-23}$ | $7.6 \times 10^{-21}$ |
| DNA Replication | 36 | 2.8 | $3.8 \times 10^{-18}$ | $9.2 \times 10^{-16}$ |
| Translational Initiation | 27 | 2.1 | $2.1 \times 10^{-17}$ | $4.7 \times 10^{-15}$ |
| U2-Type Prespliceosome Assembly | 19 | 1.5 | $5.6 \times 10^{-17}$ | $1.2 \times 10^{-14}$ |
| Ribosomal Large Subunit Assembly | 16 | 1.2 | $5.2 \times 10^{-16}$ | $1.0 \times 10^{-13}$ |

Negative Correlation (-0.4 Pearson or lower)

B

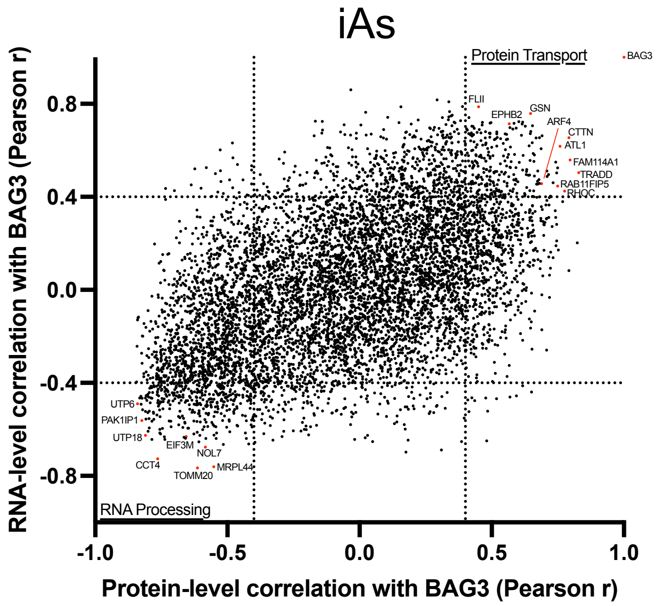


C

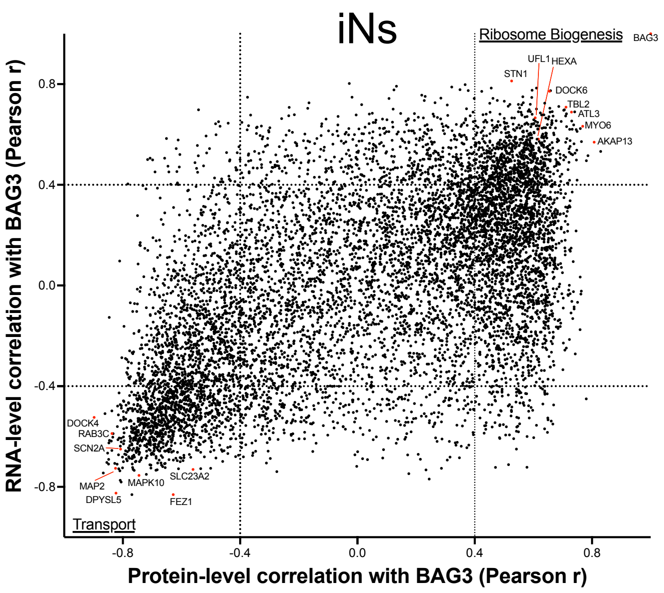


D

#### ROSMAP iNs

| Term | Count | % | P-value | Adj. P-val |
|---|---|---|---|---|
| Ribosome Biogenesis | 26 | 3.2 | $9.6 \times 10^{-13}$ | $1.0 \times 10^{-10}$ |
| rRNA Processing | 21 | 2.6 | $2.0 \times 10^{-8}$ | $1.1 \times 10^{-6}$ |
| Cell Division | 44 | 5.5 | $9.1 \times 10^{-8}$ | $3.2 \times 10^{-6}$ |
| Mitosis | 33 | 4.1 | $1.1 \times 10^{-6}$ | $2.9 \times 10^{-5}$ |
| Protein Biosynthesis | 22 | 2.7 | $2.4 \times 10^{-6}$ | $5.0 \times 10^{-5}$ |
| Cell Cycle | 54 | 6.7 | $2.3 \times 10^{-5}$ | $4.1 \times 10^{-4}$ |
| mRNA Splicing | 31 | 3.9 | $3.4 \times 10^{-5}$ | $5.1 \times 10^{-4}$ |
| mRNA Processing | 35 | 4.4 | $1.4 \times 10^{-4}$ | $1.8 \times 10^{-3}$ |
| Amino-Acid Biosynthesis | 7 | 0.9 | $1.0 \times 10^{-3}$ | $1.1 \times 10^{-2}$ |
| Translation Regulation | 15 | 1.9 | $2.2 \times 10^{-3}$ | $1.9 \times 10^{-2}$ |

Positive Correlation (+0.4 Pearson or higher)

| Term | Count | % | p-value | Adj. p-value |
|---|---|---|---|---|
| Transport | 176 | 18.1 | $9.6 \times 10^{-19}$ | $1.1 \times 10^{-16}$ |
| Neurogenesis | 53 | 5.5 | $2.0 \times 10^{-18}$ | $1.1 \times 10^{-16}$ |
| Protein Transport | 77 | 7.9 | $2.3 \times 10^{-15}$ | $8.3 \times 10^{-14}$ |
| Exocytosis | 22 | 2.3 | $9.2 \times 10^{-13}$ | $2.5 \times 10^{-11}$ |
| Endocytosis | 24 | 2.5 | $9.3 \times 10^{-9}$ | $2.0 \times 10^{-7}$ |
| Lipid Biosynthesis | 25 | 2.6 | $6.9 \times 10^{-7}$ | $1.3 \times 10^{-5}$ |
| Cholesterol Biosynthesis | 9 | 0.9 | $5.5 \times 10^{-6}$ | $8.5 \times 10^{-5}$ |
| Cell Adhesion | 41 | 4.2 | $1.4 \times 10^{-4}$ | $1.7 \times 10^{-3}$ |
| Lipid Metabolism | 55 | 5.7 | $3.2 \times 10^{-4}$ | $3.1 \times 10^{-3}$ |
| Ion Transport | 46 | 4.7 | $2.1 \times 10^{-3}$ | $1.6 \times 10^{-2}$ |

Negative Correlation (-0.4 Pearson or lower)

E

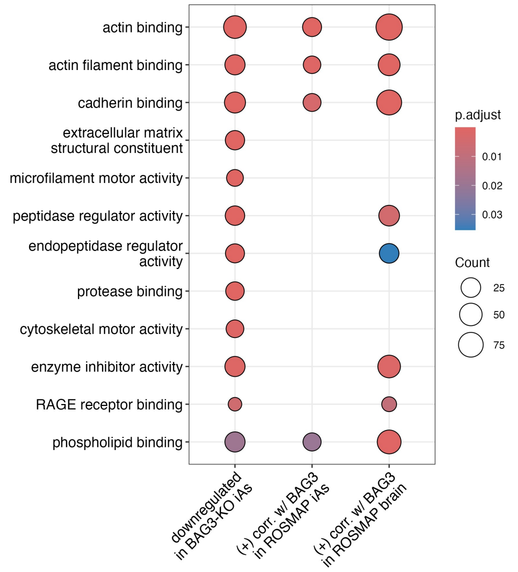


F

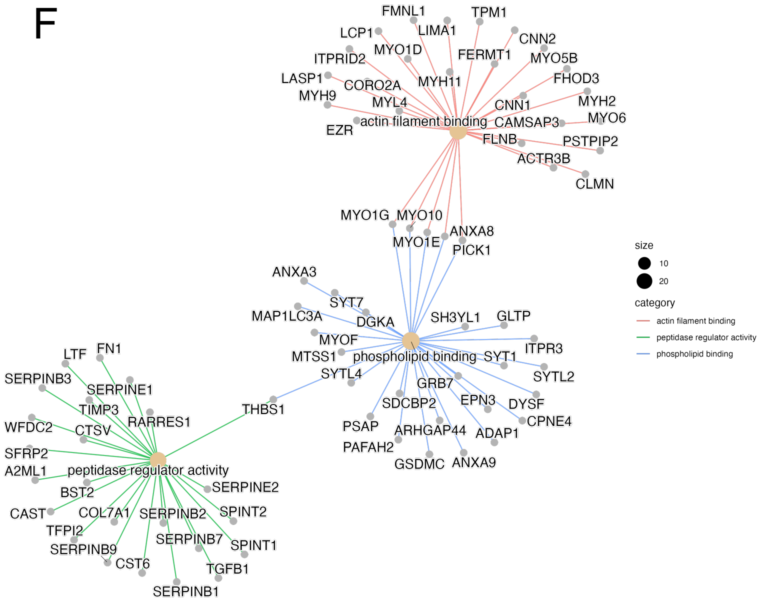

### Supplementary Fig. S3

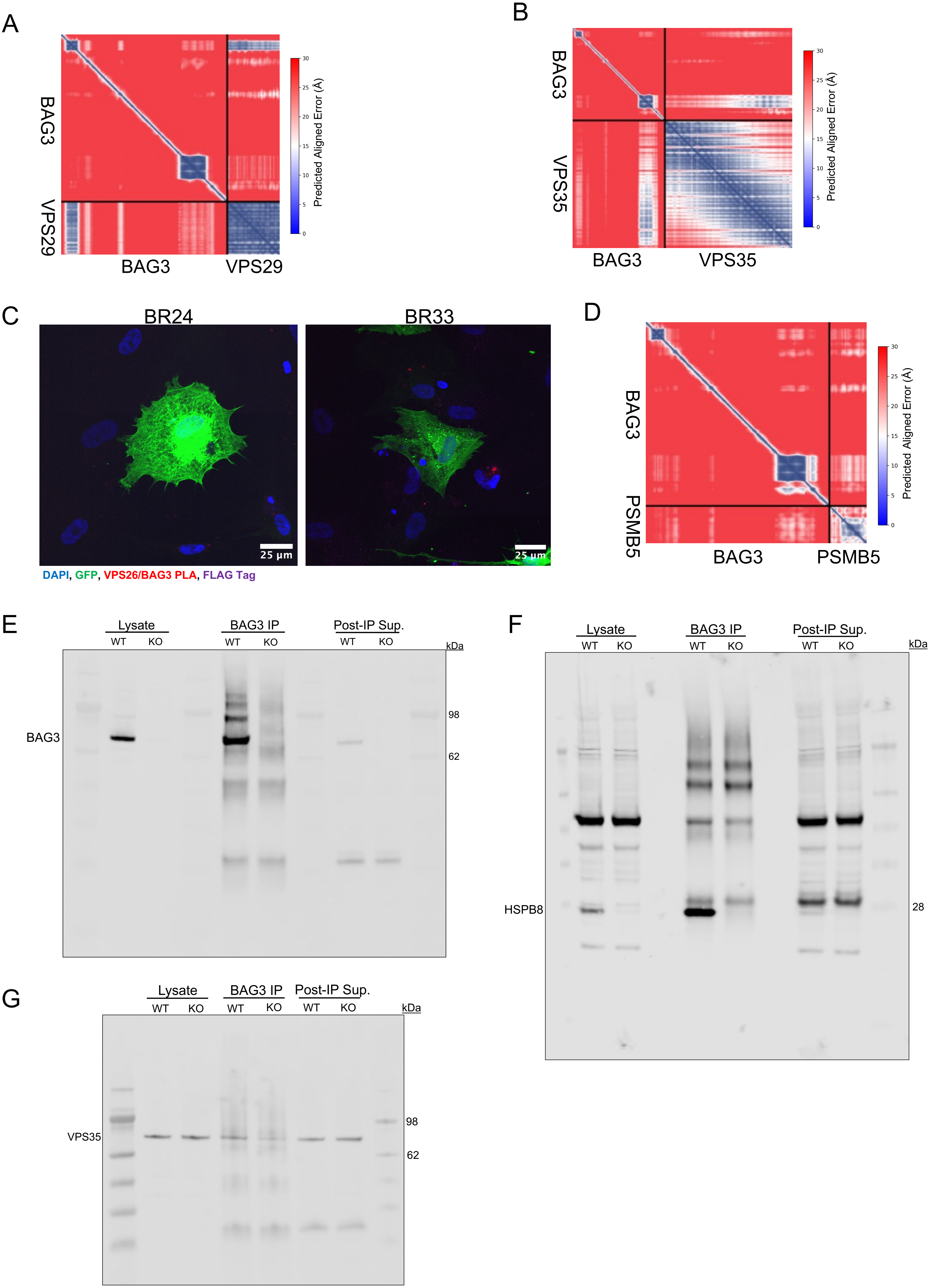

### Supplementary Fig. S4

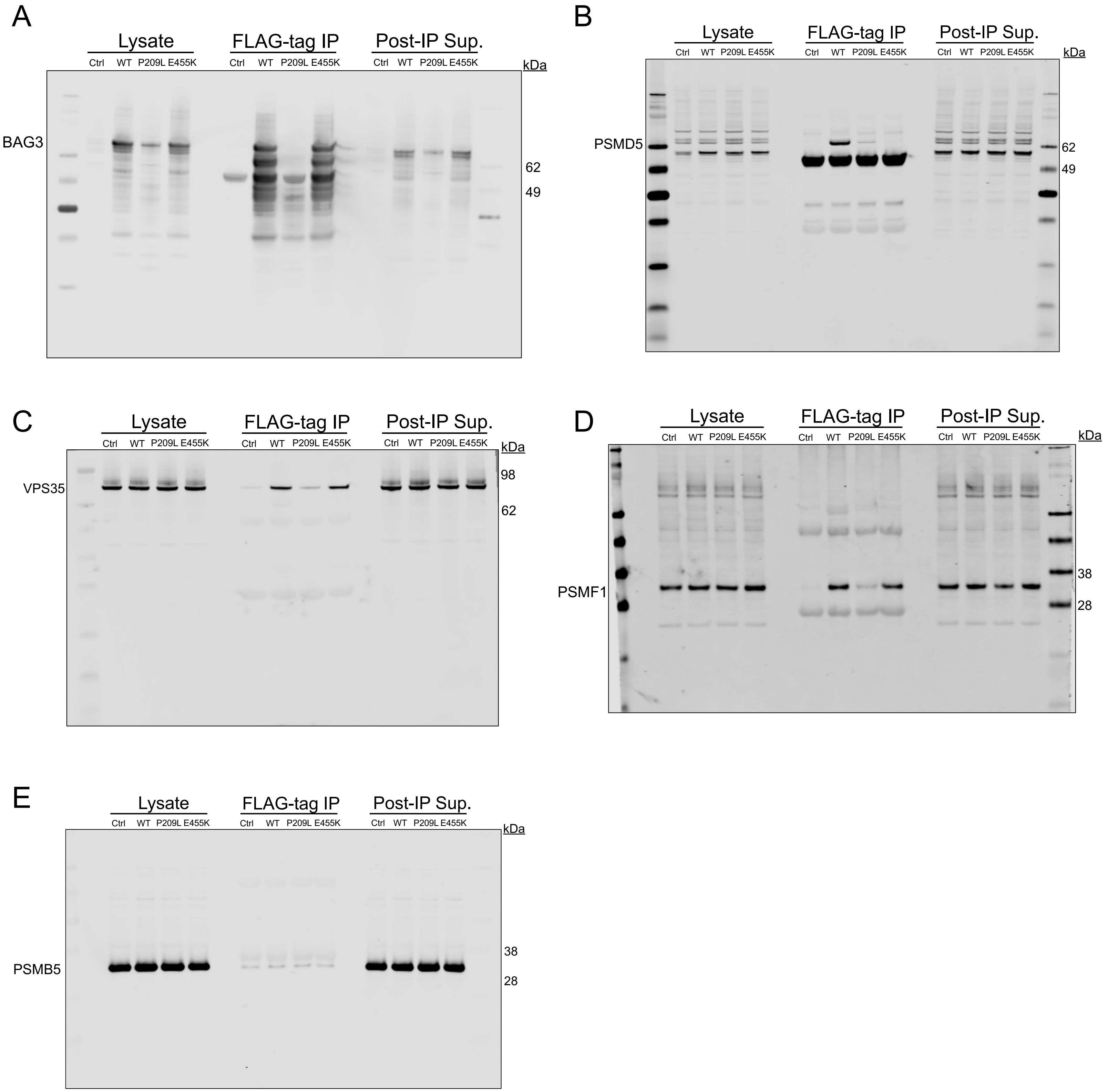

Supplementary Fig. S5

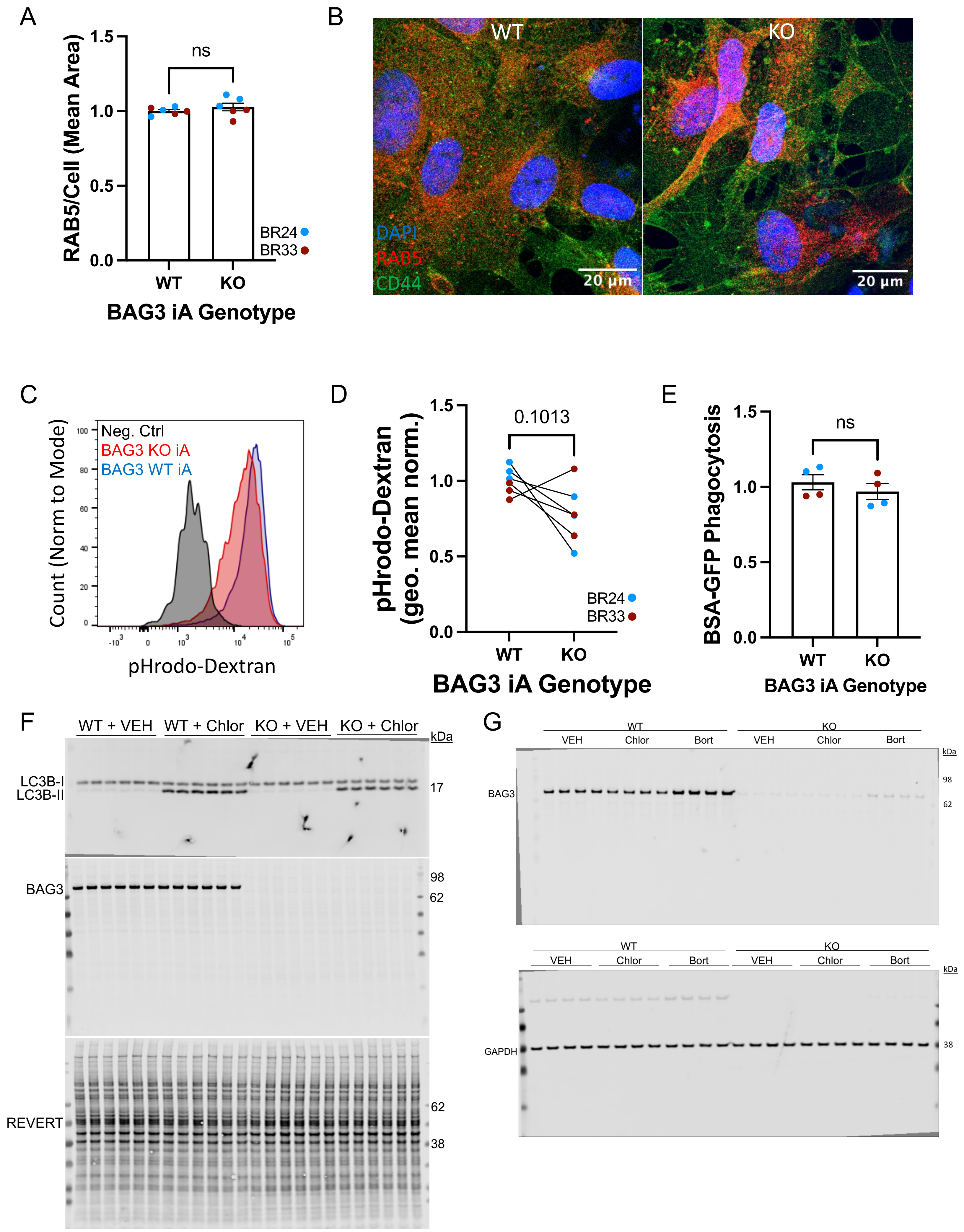

### Supplementary Fig. S6

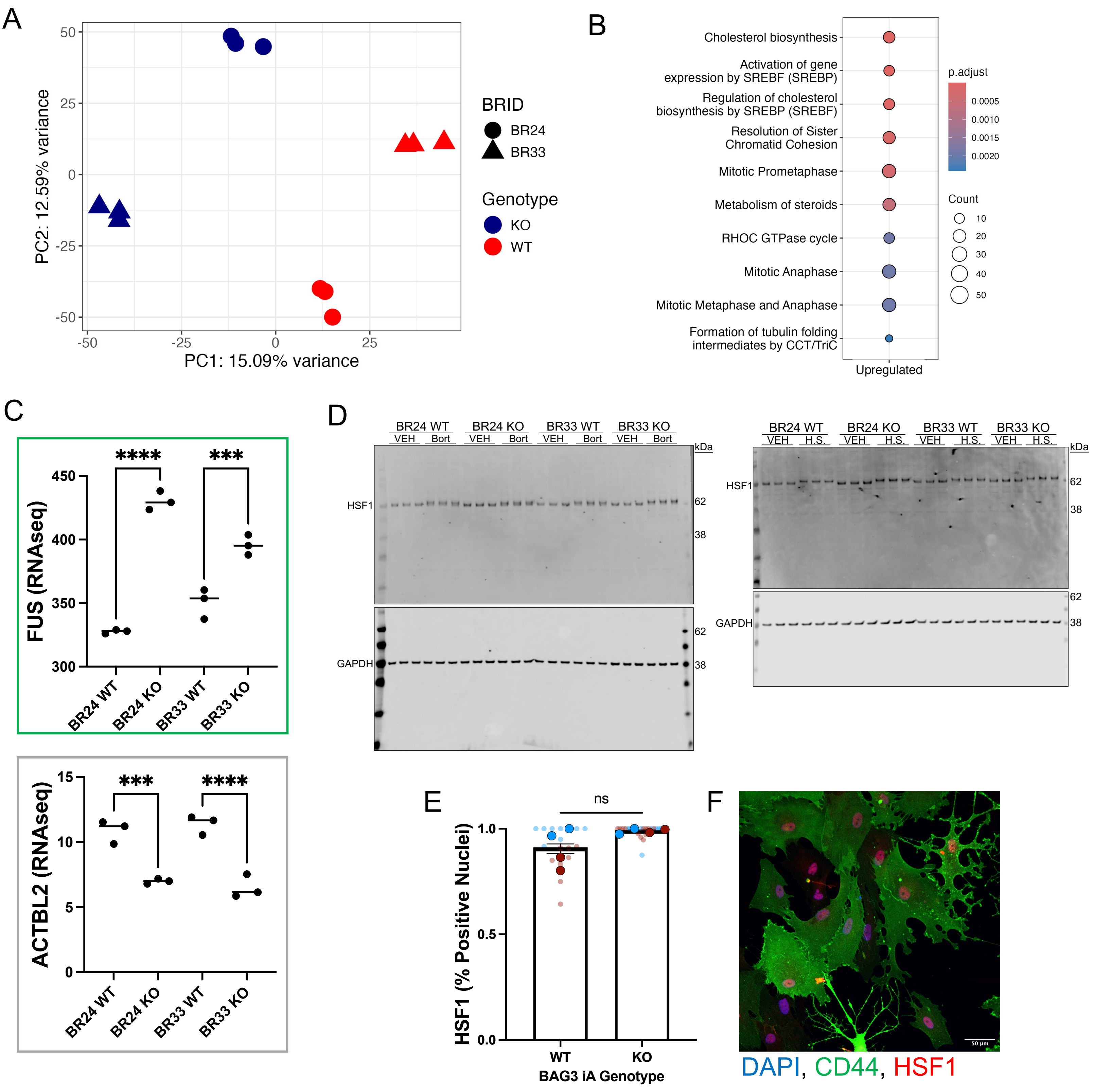

### Supplementary Fig. S7

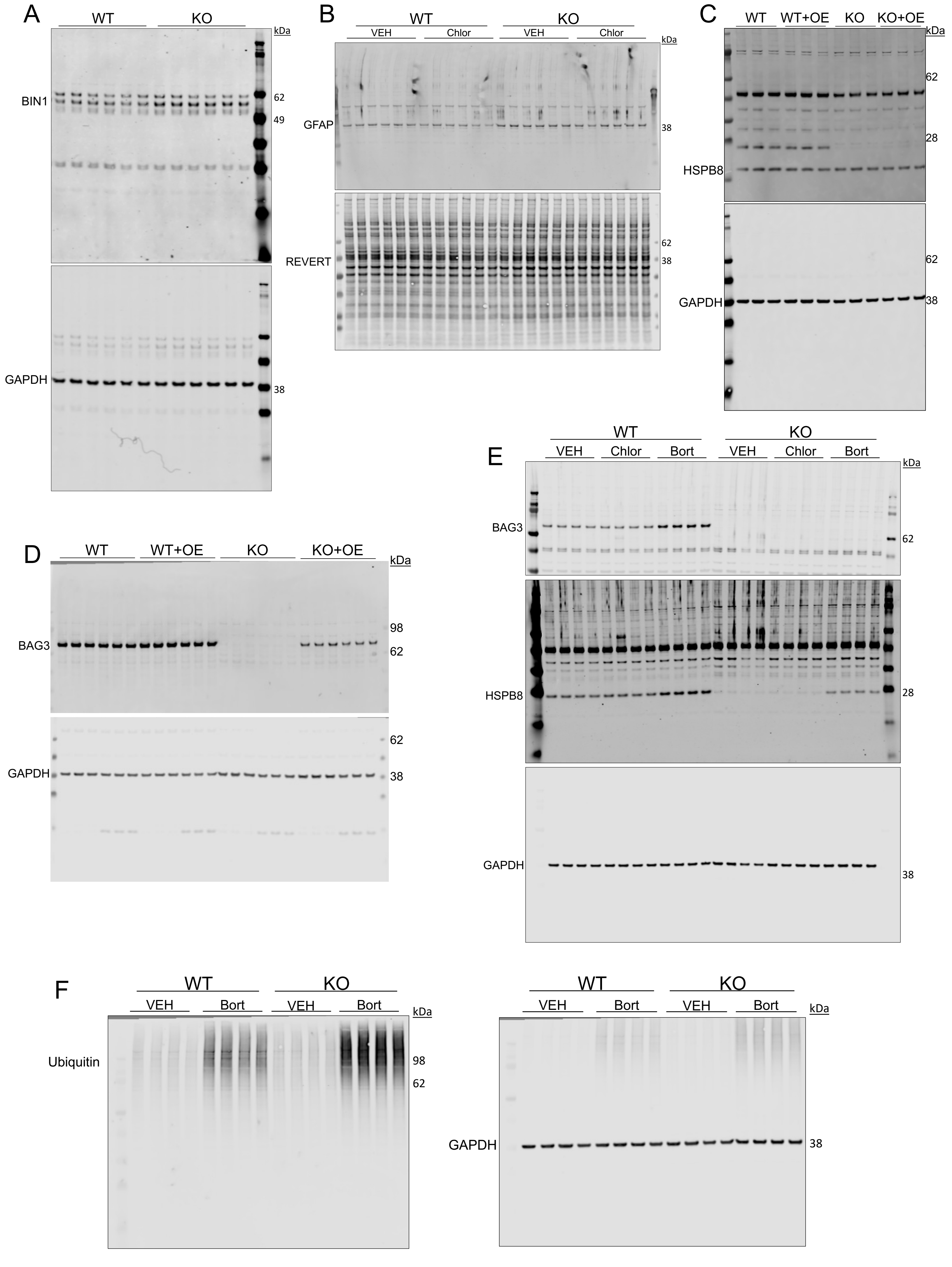

Supplementary Fig. S8

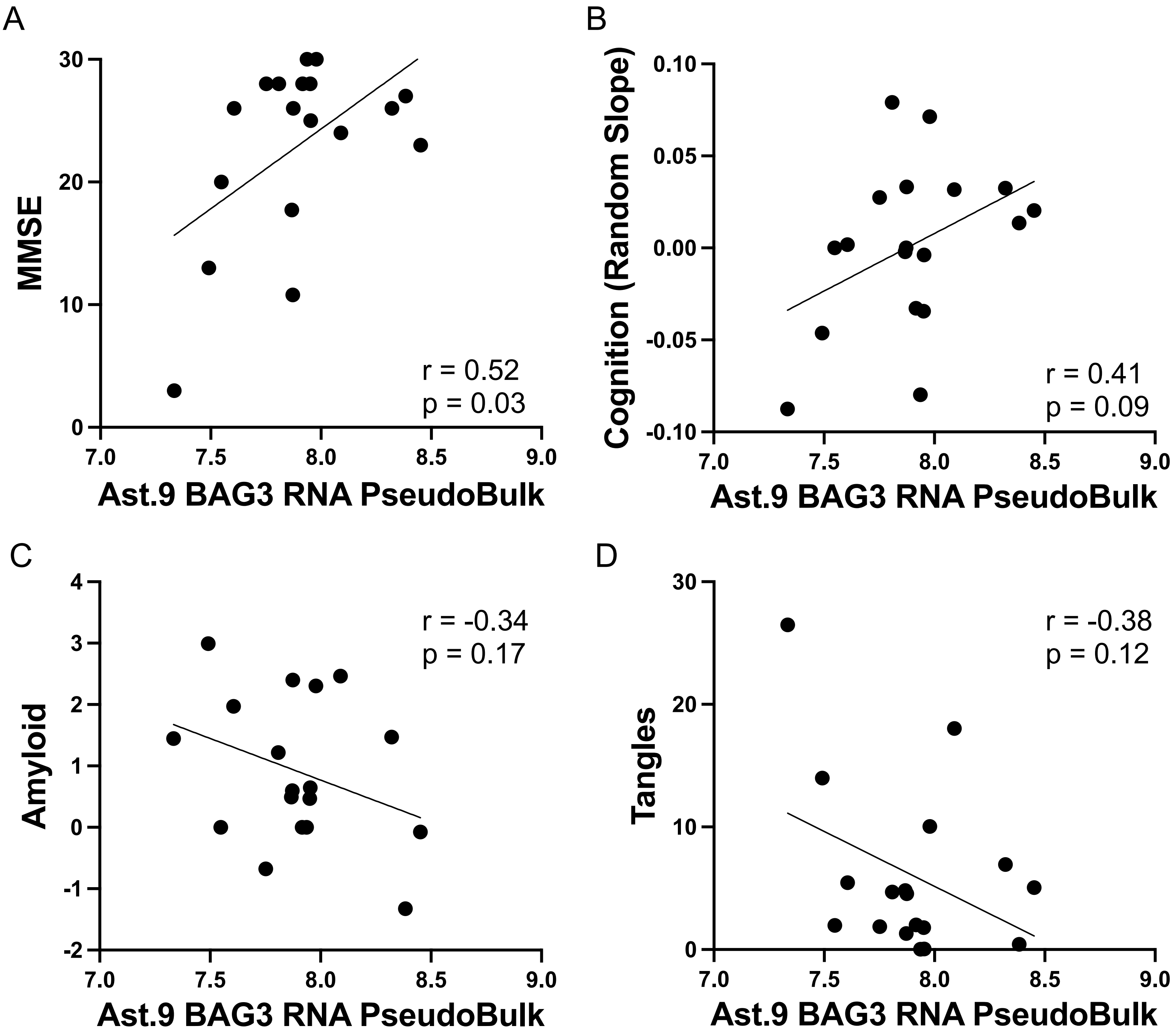
